# The kinesin Kip2 promotes microtubule growth using a bipartite polymerase module to deliver tubulin to microtubule plus-ends

**DOI:** 10.1101/2024.07.19.604271

**Authors:** Simos Nadalis, Aymeric Neyret, Ariane Abrieu, Hauke Drechsler, Dimitris Liakopoulos, Didier Portran

## Abstract

Kinesin molecular motors are essential for fundamental cellular processes such as chromosome segregation or vesicular transport. To fulfil their function, some kinesins promote microtubule growth, but the molecular mechanism underlying this activity remains unclear. One of the motors with the strongest microtubule growth-promoting activity is Kip2, a kinesin that is required for astral microtubule integrity and spindle positioning in yeast. Here we show that the ability of Kip2 to polymerize microtubules is coupled to binding and transport of free tubulin. We report that the Kip2 N-terminus is required to promote microtubule elongation *in vitro* and *in vivo*. Kip2 binds free tubulin through this unstructured, basic domain and delivers it to microtubule plus-ends. In addition to the N-terminus, we find that ATP hydrolysis and motor activity is also required for microtubule polymerisation. Finally, transfer of the Kip2 N-terminus to kinesin-1, a kinesin that lacks polymerase activity, transforms kinesin-1 into a tubulin-transporting microtubule polymerase. We propose that motor-driven tubulin delivery to microtubule plus-ends is an efficient mechanism used by kinesins to promote microtubule polymerization.

## INTRODUCTION

Kinesin motors often possess a unique combination of properties: motility and ability to control microtubule (MT) dynamics (Wu *et al*, 2006; Hunter & Wordeman, 2000; Su *et al*, 2012). In this manner, kinesins coordinate their movement with MT growth and shrinkage i.e. during chromosome segregation, cargo transport or cell polarization. A large number of studies has been dedicated to molecular motors of the kinesin-8 and kinesin-13 families that act as MT-depolymerases (Friel & Welburn, 2018; Shrestha *et al*, 2018). In contrast, only few examples of MT-polymerizing kinesins have been studied. These include the kinesin-5 Eg5 (Chen *et al*, 2019), and the kinesin-7 CENP-E (Sardar *et al*, 2010).

The kinesin with probably the strongest MT-growth-promoting activity is the plus-end-directed kinesin Kip2 of budding yeast. Homologous to the fission yeast Tea2 (Browning *et al*, 2000), Kip2 is a central regulator of yeast astral microtubules, termed cytoplasmic MTs (cMTs). Similar to *tea2Δ* null cells, cells lacking Kip2 display extremely short cMTs, whereas Kip2 overexpression results in abnormal cMT elongation (Cottingham & Hoyt, 1997; Carvalho *et al*, 2004). In addition to regulating cMT length, Kip2 transports the spindle positioning factors dynein and Kar9 to the plus-ends of cMTs (Lee *et al*, 2005; Maekawa *et al*, 2003; Markus *et al*, 2009). Consequently, Kip2 is required for correct orientation of the mitotic spindle relative to the cleavage apparatus, a process essential for chromosome segregation in yeast (Miller *et al*, 1998; Cottingham & Hoyt, 1997).

Kip2 forms complexes with the MT-associated proteins Bim1/EB1 and Bik1/CLIP-170 that act as accessory factors, increasing processivity and residence time of the motor at MT plus-ends, respectively (Drechsler *et al*, 2015; Roberts *et al*, 2014; Chen *et al*, 2023). Both proteins bind to a N-terminal stretch of app. 100 amino acids preceding the motor domain of Kip2. This highly basic and very likely unstructured region is targeted by Cdc28/Cdk1, Mck1/GSK-3 and Dbf2 kinases, that control the interaction of Kip2 with its accessory factors as well as its microtubule substrate through phosphorylation (Drechsler *et al*, 2015). *In vitro* characterization of Kip2 showed that it directly promotes MT growth by increasing the rate of tubulin association *k_on_* and concomitantly decreasing the tubulin dissociation rate *k_off_*, thereby decreasing the frequency of catastrophes almost 20-fold (Hibbel *et al*, 2015). In addition, Kip2 increases the speed of MT polymerization *in vitro* by 3-fold, without the need of accessory factors (Hibbel *et al*, 2015; Bowne-Anderson *et al*, 2015). Hibbel *et al*., proposed that Kip2 acts either as a processive MT polymerase or as a shuttle that transports tubulin to MT plus-ends. Indeed, another recent study showed that Kip2 interacts with free tubulin via an extra site (the P1 site) in its motor domain (Chen *et al*, 2023). Mutation at this site severely reduced the growth promoting activity of Kip2, but it is unclear how this site could participate in tubulin shuttling, since it is found in the MT-binding motor domain.

Here, we present evidence that Kip2 uses its N-terminal domain to bind free tubulin dimers and shuttle them to MT plus-ends. Deletion of the Kip2 N-terminus abrogates tubulin transport as well as the ability of Kip2 to promote MT growth *in vivo* and *in vitro*. Importantly, the motor activity of Kip2 is required for its MT-growth-promoting activity. Fusing the Kip2 N-terminus to kinesin-1 enables kinesin-1 to transport tubulin and to promote MT growth *in vitro*. Together, this data suggests that Kip2 acts as MT polymerase that paves its own way (Steinberg, 2007) by transporting and adding tubulin dimers to MT plus-ends.

## RESULTS

### 1. The Kip2 polymerase activity depends on the N-terminus of the kinesin

In search for the molecular mechanism used by Kip2 to promote cMT growth, we characterized the biophysical properties of different recombinant Kip2 constructs (Fig. 1A) in *in vitro* reconstitution assays. A short, “bonsai” bKip2-mCherry version (bKip2-WT) that comprises the motor domain and the first putative coiled-coil dimerization domain up to amino acid 560, purified from bacteria (Fig. S1A) has been shown to recapitulate the properties of full length Kip2 ((Chen *et al*, 2023; Hibbel *et al*, 2015), Fig. 1B and Fig. S1C, E, F, purified proteins Fig. S1A, B). Indeed, the bKip2-WT version accumulated at the tips of growing MTs (Fig. S1C), strongly suppressed catastrophes and increased MT polymerization speed in a concentration-dependent manner, similar to full length Kip2 (Fig.1B-D). Moreover, bKip2-WT was able to polymerize MTs at tubulin concentrations as low as 0.5µM (Fig. S1C) and accelerated tubulin polymerization, similar to full length Kip2 ((Fig. S1D), (Hibbel *et al*, 2015)). Finally, bKip2-WT moved processively on reconstituted GMP-CPP MTs (Fig. S1E-G)) with similar velocity to full length Kip2 ((Fig. S1G and S2H), 8.0+/-2.1 µm/min for bKip2-WT and 6.1+/-1.5 µm/min for Kip2-WT). This data shows that the C-terminal part of Kip2 (aa 561-706) is dispensable for its MT-growth-promoting activity *in vitro*. Bonsai bKip2-WT versions were thus used for all following experiments, unless otherwise indicated.

**Figure 1:**
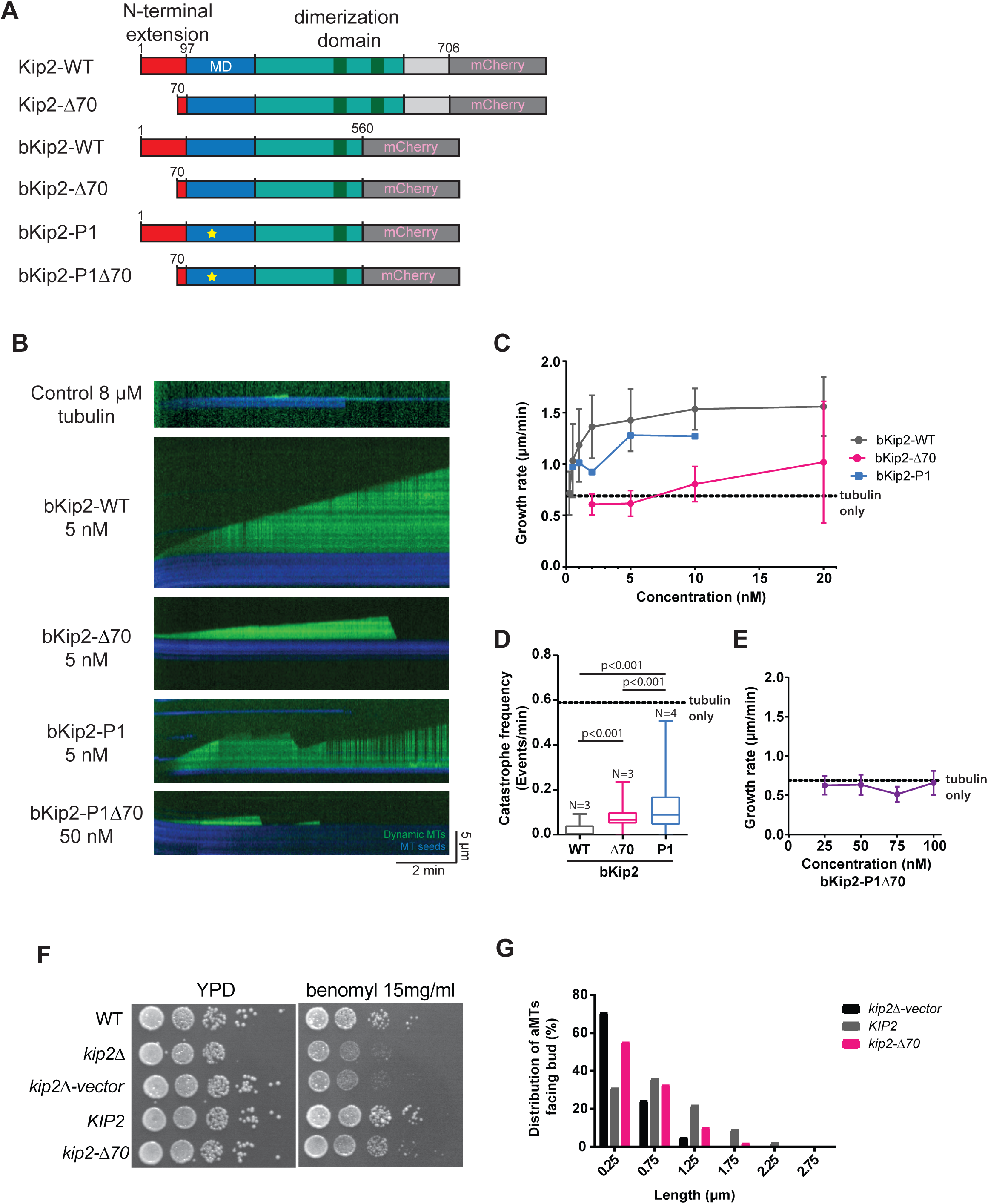
The N-terminus of Kip2 is required for microtubule stabilization *in vitro* and in i*n vivo*. A) Scheme of the different constructs used in this article. Red: unstructured N-terminal region of Kip2, blue: motor domain (MD), light green: dimerization domain dark green: coiled-coiled domain, dark gray: C-terminal mCherry tag. The yellow star represents the interface 1 mutation on the motor domain, named P1 (Chen *et al*., 2023). B) Kymographs extracted from TIRFm images of dynamic MT assays using the indicated Kip2 versions at 5nM concentration. Blue: MT seeds, green: atto-488-tubulin C) MT growth rate as a function of kinesin concentration in presence of different bKip2 mutants in presence of 8 µM tubulin (mean ± SD of mean). The dashed line shows the MT growth rate at 8 µM tubulin concentration in absence of kinesin. Experiments were repeated at least 3 times, except for the control that was repeated 2 times. D) Quantification of catastrophe frequencies in presence of different bKip2 mutants at 5nM concentration. The bar represents the median, the box marks the interquartile range, and the vertical line covers the 95% confidence interval. The dashed line shows the control: only tubulin. statistical test: Mann-Whitney. E) MT growth rate as a function of kinesin concentration for bKip2-P1Δ70 mutants (mean ± SD of mean), the dashed line shows the MT growth rate at 8 µM tubulin concentration in absence of kinesin. F) Benomyl sensitivity assay. Shown are serial dilutions of yeast mutant cells of the indicated genotype grown on full medium plates (YPD) without and with 15 mg/ml benomyl. G) Length distribution of cMTs emerging from bud-oriented SPBs for the mutants indicated. Lengths were measured from maximum intensity projection images of GFP-Tub1-expressing cells.

Kip2 promotes MT growth by increasing the rate of MT growth and also by acting as an anti-catastrophe factor (Hibbel *et al*, 2015; Bowne-Anderson *et al*, 2015). To find which domains of Kip2 are important for these activities, we quantified the MT growth and catastrophe rates for three different bKip2 constructs: bKip2-Δ37 that lacks the Bim1/EB1 binding site, bKip2-Δ70 that lacks a large part of the phospho-regulated N-terminus (Drechsler *et al*, 2015), and the motor domain P1 mutant (K294R K296R, (bKip2-P1), lacking a tubulin binding site at the motor domain (Chen *et al*, 2023) (Fig. 1A, purified proteins Fig. S1A, B). The bKip2-Δ37 construct behaved essentially like wild-type for all parameters tested (Fig. S1J, K). Measuring the fluorescence intensity at different concentrations of the kinesin on Taxotere-stabilized MTs, we found that the affinities of bKip2-Δ70 and bKip2-P1 were 1.3 times and 3.5 lower compared to bKip2-WT, respectively (Fig. S1I). Both mutations caused a decrease in the anti-catastrophe activity of the kinesin (Fig. 1D).

The bKip2-P1 mutant displayed reduced MT growth rates for all concentrations tested (Fig. 1C). The most striking effect however was observed for bKip2-Δ70 that was severely defective in MT growth and lost its ability to increase the rate of MT growth over a large range of concentrations (Fig. 1B, C). Analysis of full-length Kip2-Δ70 purified from insect cells confirmed these results (Fig. S2A-C). To examine whether the Kip2 N-terminus is important for maintenance of cMT length also *in vivo*, we expressed untagged Kip2-Δ70 at endogenous levels in yeast cells (Fig. S1L). Kip2-Δ70-expressing cells displayed clearly shorter MTs, with a length distribution falling between the *kip2Δ* null and wild-type cells and, in line with this phenotype, a corresponding benomyl sensitivity to the microtubule-destabilising drug benomyl (Fig. 1F, G). Together, these results clearly suggest that the polymerase activity of Kip2 depends mainly on its unstructured, N-terminal domain, and also on its motor domain, as already described (Chen *et al,* 2023). Indeed, the double bKip2-P1Δ70 mutant was unable to increase the rate of MT polymerization or to promote MT growth at any concentration up to 100nM (Fig. 1E).

### 2. Kip2 transports free tubulin dimers to microtubule plus-ends via its N-terminus

Two models have been proposed to explain how Kip2 might promote the growth of MTs (Hibbel *et al*, 2015): either Kip2 promotes the addition of free tubulin subunits at MT plus-ends, or alternatively, it acts as a shuttle that transports free tubulin to the plus-ends of MTs, thereby increasing tubulin concentration locally to promote MT polymerization. In favour of the shuttle model, Kip2 was recently shown to bind free tubulin in solution via a positively charged motif at the P1 site in its motor domain (Chen *et al*, 2023).

We thus investigated whether Kip2 could transport tubulin dimers to MT plus-ends and performed motility assays at different kinesin concentrations (4-20nM) on GMP-CPP stabilized MTs, in presence of 50nM free, ATTO-488-labelled tubulin. Indeed, we observed tracks of green tubulin moving along stabilized MTs (Fig. 2A and Movie 1). These tracks overlapped with tracks of bKip2-WT, whereas their number increased with increasing concentration of the kinesin (Fig. 2B).

**Figure 2:**
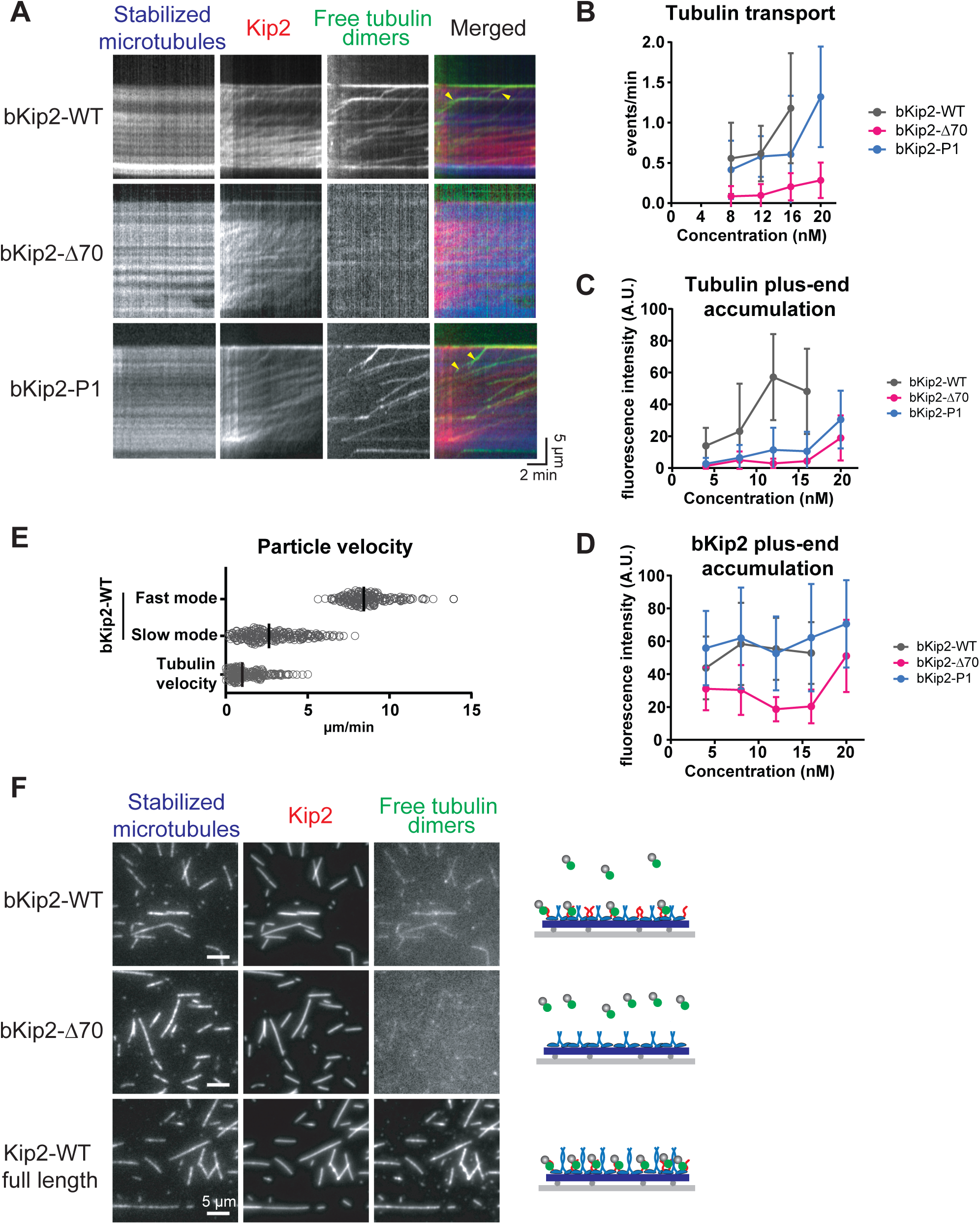
Kip2 captures and transports free tubulin to microtubule plus-ends via its N-terminus. A) Kymographs extracted from motility assays of bonsaï bKip2 mutants (16 nM for bKip2-WT, 20 nM bKip2-P1 and 20 nM bKip2-Δ70) on stabilized MTs spiked with with free atto-488 labeled-tubulin (50nM). Blue: MT seeds, red: kinesin and green: atto-488-tubulin. Note the green tubulin traces that colocalize with kinesin, that are absent in the bKip2-Δ70 mutant. B) Quantification of the frequency of tubulin dimer transportation events per min, per MT, at different concentrations of bKip2 constructs (mean ± SD of mean). C) Quantification of plus-end accumulation of tubulin in the different kinesin mutants indicated (mean ± SD of mean). D) Quantification of plus-end accumulation of the kinesin mutants indicated (at least 52 MTs were analyzed per condition) (mean ± SD of mean). E) Quantification of the velocity of bKip2-WT (at 5 nM) at fast mode (single motors) and slow mode (clusters) and of the velocity of tubulin particles at 8 nM of bKip2-WT and 50 nM of atto-488 labeled-tubulin (Each circle represents a velocity measurement, the solid bar represents the median). F) TIRFm images of bKip2, bKip2-Δ70 and full-length Kip2-WT at 5nM, attached with AMP-PNP on GMP-CPP-stabilized MTs, followed by addition of free atto-488 labeled-tubulin (50nM). Blue: MTs, red: kinesin and green: atto-488-tubulin. A model of the assay and the result is shown on the right of the images. Scale bar = 5 µm.

Importantly, deletion of the Kip2 N-terminus drastically decreased the number of transport events that were almost absent at concentrations of 8 and 12nM of bKip2-Δ70 (Fig. 2B). Very similar results were obtained when we used full-length Kip2 versions (Fig. S2D-F). Here also, the occurrence of tubulin transport events by Kip2-Δ70 was reduced 3- to 8-fold compared to wild type Kip2, depending on the kinesin concentration (Fig. S2F). The residual small number of transport events by bKip2-Δ70 or its full-length counterpart was mainly due to occasional transport of large tubulin clusters (Fig. S2E). In summary, these experiments suggest that Kip2 transports free tubulin in vitro, and that the N-terminus of the kinesin is required for the transport process.

Tubulin was not only transported along stabilized MTs, but also accumulated together with bKip2-WT at their plus-ends (Fig. 2A, C and Fig. S2D). Both the amount of tubulin dimers and bKip2-WT present at the ends of MTs increased along with the concentration of bKip2-WT in the assay (Fig. 2C, D, and Fig. S2I, J). In contrast, kinesin and tubulin accumulation at plus-ends was severely reduced in absence of the first 70 N-terminal amino acids of Kip2 (Fig. 2A, C, D). We thus concluded that the N-terminus of Kip2 is required not only for tubulin transport, but also for retention of Kip2 and tubulin at MT plus-ends.

Since the P1 site in the motor domain binds tubulin dimers and is required for the MT-growth-promoting activity of Kip2 (Chen *et al*, 2023), we subsequently tested whether the P1 site in the motor domain is also required for tubulin transport. We observed that tubulin transport by bKip2-P1 was slightly reduced in the assay (Fig. 2B). Accumulation of bKip2-P1 at MT plus-ends was not affected (Fig. 2D), despite bKip2-P1 having a lower affinity for MTs than bKip2 (Fig. S1G). Remarkably, tubulin accumulation at MT plus-ends by bKip2-P1 was reduced compared to bKip2-WT (Fig. 2C).

These results show that the N-terminal amino acid stretch of Kip2, but not the P1 site, is required for transport of free tubulin along MTs. In contrast, accumulation of tubulin at MT plus-ends requires both the Kip2 N-terminus and the P1 site.

Lastly, we examined whether the Kip2 N-terminus is required to bind tubulin. For this, we decorated stabilized MTs with either bKip2-WT or bKip2-Δ70 and subsequently added fluorescent free tubulin in presence of non-hydrolyzable AMP-PNP to avoid kinesin movement (Fig. 2F). Free tubulin bound to bKip2-WT but not to bKip2-Δ70-decorated microtubules, suggesting that the Kip2 N-terminus indeed interacts with tubulin directly. When full-length Kip2-mCherry from insect cells was used, affinity for tubulin was even higher, implying that the full-length kinesin binds tubulin in a cooperative fashion (Fig. 2F). In view of these results, we concluded that Kip2 binds tubulin and transports it to MT plus-ends via its N-terminus.

### 3. Kip2 can transport tubulin in different modes

We observed that Kip2 was able to form larger assemblies or “clusters” on MTs (Fig. S5A, B). Cluster formation depended on Kip2 concentration, the nature of the MT lattice and correlated with slower movement on MTs. To better visualize this behaviour, we generated MTs that consisted of one Taxotere-stabilized GDP segment and a second, GMPCPP segment. Indeed, bKip2-WT moved on the GDP lattice mostly as clusters of low speed (Fig. S5A-C), median velocity: 2.6+/-1.7 µm/min). These would then separate into fast particles after entering the GMPCPP segment (Fig. S5 A-C, median velocity: 8.4+/-1.5 µm/min). We could also observe this behaviour on dynamic MTs elongating from GMPCPP MT-stabilized seeds, where bKip2-WT moved more often as fast, single particles on the GMPCPP seed and as slower clusters on the dynamic MT lattice (Fig. 2E and Fig. S5D). bKip2-Δ70 behaved essentially like bKip2-WT regarding motility, however bKip2-P1 was less prone to cluster and would stay in the fast-moving mode (Fig. S5D). This difference in behaviour could be due to the lower affinity of bKip2-P1 for Taxotere-stabilized MTs (Fig.S1I) as we could observe slow clusters of bKip2-P1 on GDP-MT lattice at higher concentrations (16nM, Fig. 2A). These data suggest that the motor domain of Kip2 is sensitive to the state of the MT lattice.

To investigate, whether tubulin is transported by single Kip2 motors or by Kip2 clusters, we measured the velocity of tubulin transport events. For bKip2-WT at concentrations 8 to 16nM used in the tubulin transport assays, the velocity distribution of tubulin transport tracks indicated that most tubulin moves at a speed similar to slow-moving bKip2-WT clusters (Fig. 2E). However, we could also observe fast tubulin transport events at low concentrations of Kip2 (1nM) that allow single molecule imaging (Fig. S2D). These data suggest that both single motors as well as clusters of motors can transport tubulin.

The concentration of free tubulin is very low in the transport assays described above (50 nM). We next examined tubulin transport under kinesin and tubulin saturating conditions. We thus first immobilized Kip2 full length on MTs in presence of slowly hydrolyzable AMP-PNP, allowed fluorescent tubulin (1µM) to bind to Kip2 and subsequently washed away the fluorescent tubulin and supplied ATP to restore kinesin motility. After a lag phase, Kip2-tubulin complexes clearly merged into droplet-like structures of various sizes that moved processively towards MT plus-ends (Fig. S3A, B and Movie 2). Droplet formation was independent of free tubulin or the Kip2 N-terminus. Electron microscopy confirmed the presence of amorphous Kip2 assemblies on MTs in presence of ATP, but not under AMP-PNP conditions (Fig. S4).

In summary, Kip2 motility depends on the nature of the MT lattice. The kinesin moves slower on GDP MTs and forms clusters with increasing concentration. Tubulin transport occurs independently of the degree of Kip2 clustering and can acquire the form of droplets at high kinesin concentrations *in vitro*.

### 4. The motor activity of Kip2 is required for its polymerase activity

We next wished to shed more light into the molecular mechanism of MT polymerization by Kip2. Having four microtubule binding interfaces (twice the motor domain and twice the N-terminal extension), Kip2 dimers could act as microtubule polymerase in a XMAP215-like manner: Kip2 dimers would follow the growing microtubule plus-end by diffusion, bind free tubulin directly from solution and integrate tubulin into a growing protofilament. Since such polymerase activity does not require energy consumption in form of ATP hydrolysis (Brouhard *et al*, 2008), we asked whether Kip2 still can promote microtubule growth in the absence of ATP. We hence incubated dynamic MTs with low nanomolar amounts of bKip2-WT in the presence of ADP, which allows binding of the motor to the MT lattice, but prevents motility (Fig. 3A). Under these conditions, no MT growth was observed for kinesin concentrations up to 2nM. For bKip2-WT concentrations ≥ 5nM, MT growth was partly restored (Fig. 3A), but the MT growth rate did not exceed the growth rate in the absence of motor (Fig. 3B). Catastrophes, however, were still efficiently suppressed by bKip2-WT in ADP (Fig. 3C). Hence, the behaviour of bKip2-WT in ADP suggests that the full mechanochemical cycle of the motor is essential for the Kip2 polymerase activity.

**Figure 3:**
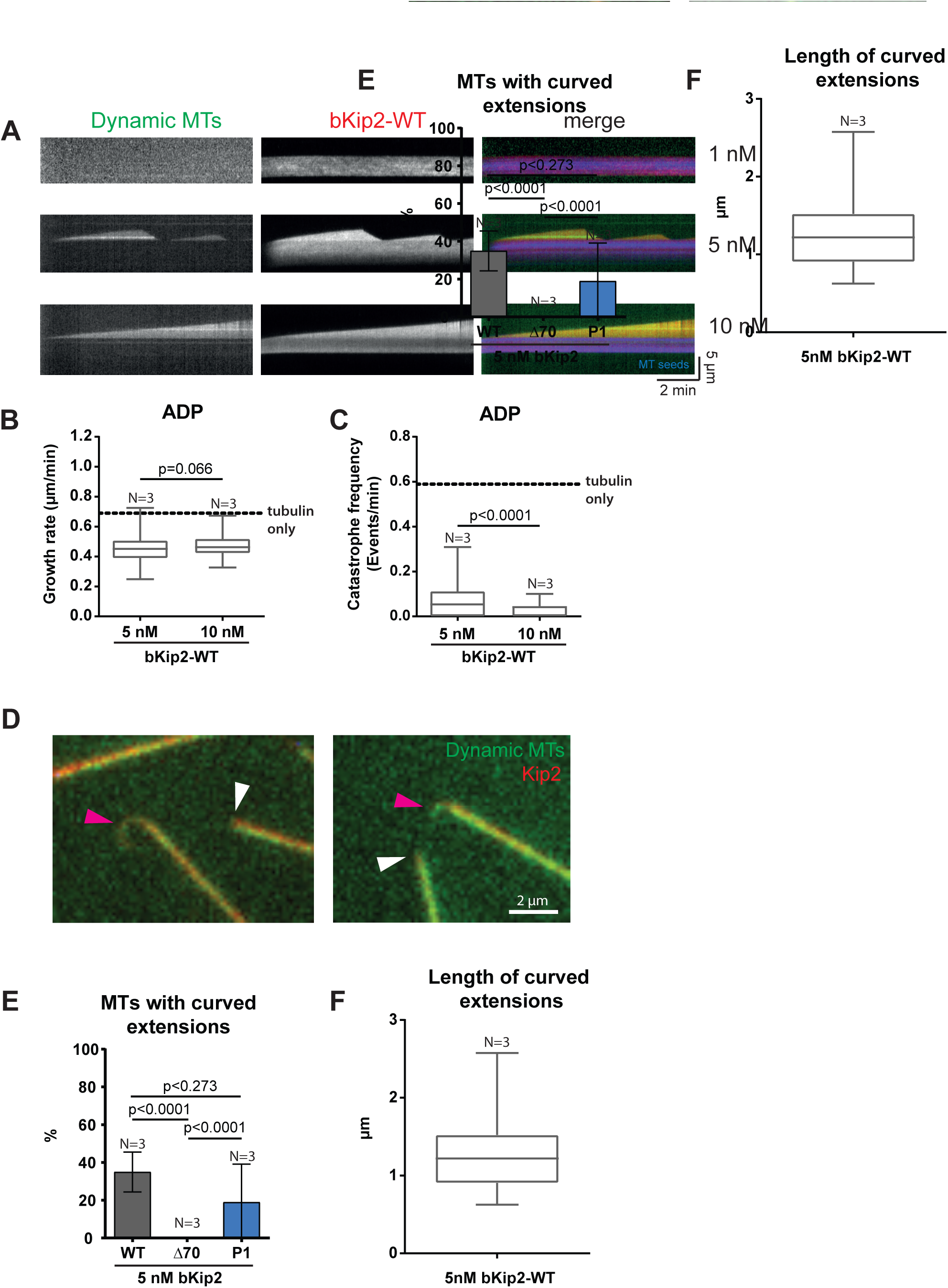
The motor activity of Kip2 is required for its polymerase activity. A) Kymographs extracted from dynamic MT assays in presence of 8 μM tubulin and bKip2 (1nM, 5nM or 10nM), ADP and GTP. L Blue: MT seeds, red: kinesin and green: atto-488-tubulin. B,C) Quantification of MT growth rate and catastrophe frequency in the assays described in (A). Control: tubulin only. D) TIRF microscopy images of curved MT extensions at the plus-ends of dynamic MTs, grown in presence of 5nM Kip2-WT. Red: kinesin and green: atto-488-tubulin. Pink arrows indicate a curved extension and the white arrow a regular growing MT plus-end. Note that both tubulin and kinesin are present on the extensions. E) Quantification of the occurrence of curved MT extensions during dynamic MT assays with bKip2, bKip2-Δ70 and bKip2-P1 (mean ± SEM). Note: no protofilament extensions were observed with bKip2-Δ70. F) Quantification of curved MT extension length with 5 nM of bKip2-WT. BFor (B), (C) and (F), the bar represents the median, the box marks the interquartile range, and the vertical line covers the 95% confidence interval. Statistical test: Mann-Whitney.

Remarkably, the plus-ends of MTs growing in presence of either bKip2-WT or bKip2-P1 often showed long, curved extensions (Fig. 3D, E), the length of which reached up to 2µm (Fig. 3F). These structures likely represented protofilaments or groups of protofilaments that were polymerized but not closed yet to a complete MT cylinder. We never observed curved extensions in presence of bKip2-Δ70 (Fig. 3E). In conjunction with the requirement of Kip2 motor activity for MT polymerization, these findings suggest that Kip2 elongates protofilaments as it walks on MTs using its N-terminus to add tubulin dimers to their plus-ends.

### 5. The Kip2 N-terminus can transform kinesin-1 to a MT polymerase

Combining motor domain motility with the tubulin-binding property of the Kip2 N-terminus seemed to create a module that promotes MT polymerization. To further test this idea, we fused the 97 N-terminal amino acids of Kip2 to an mCherry-tagged, truncated version of the kinesin-1 KIF5B (Fig. 4A), that does neither decrease the rate of catastrophes, nor alters the rate of MT polymerization (Budaitis *et al*, 2022; Andreu-Carbó *et al*, 2022) and asked, whether the ^NKip2^KIF5B chimera was capable to transport tubulin and to promote MT growth (Fig. 4B-I). Wild-type KIF5B and the chimeric ^NKip2^KIF5B had comparable velocities (Fig. S6B, C), while the affinity of ^NKip2^KIF5B for MTs was significantly higher, compared to KIF5B (Fig. S6D).

**Figure 4:**
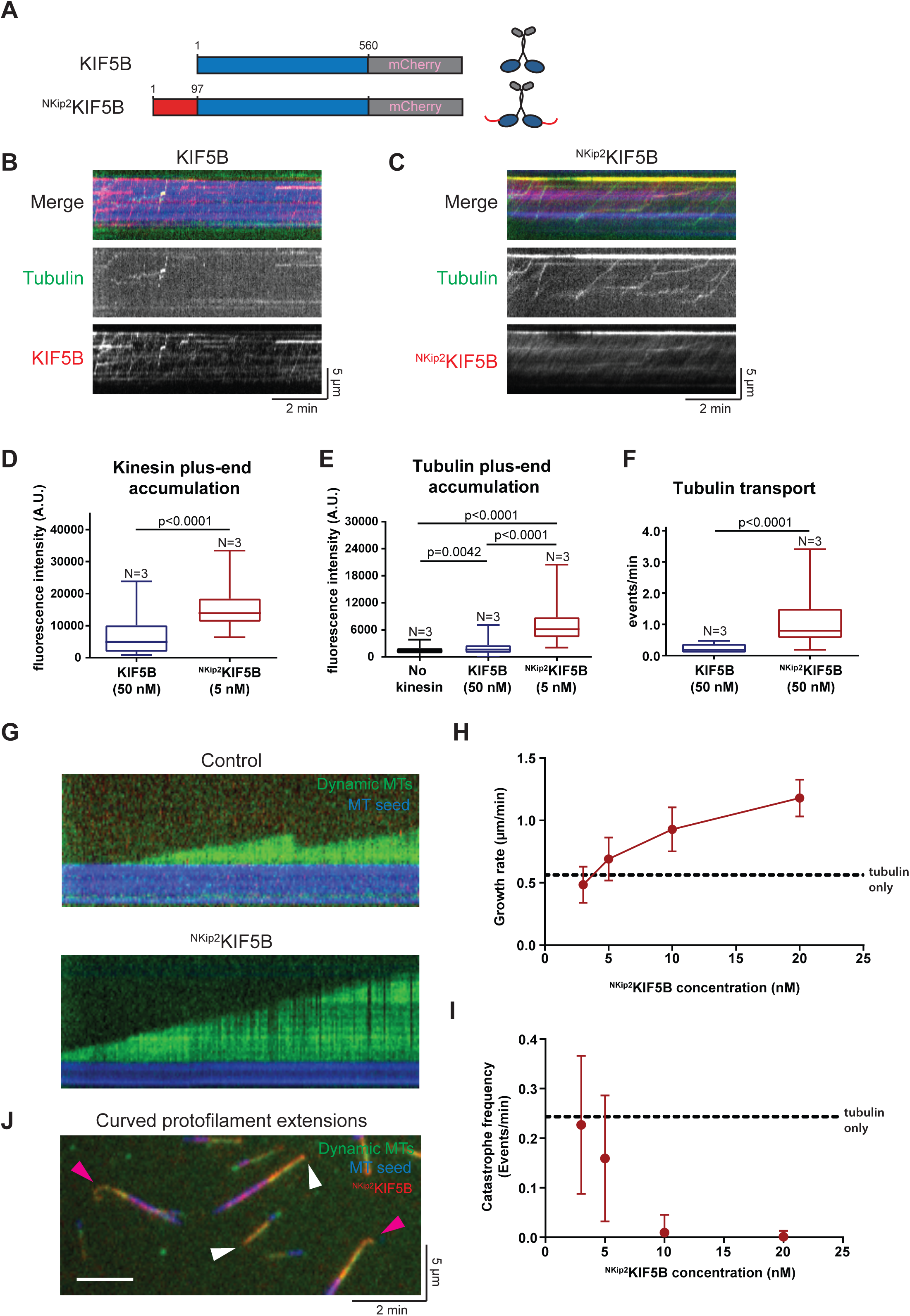
The Kip2 N-terminus is a transferable tubulin delivery domain. A) Scheme of the different constructs used in the study. In blue represent the C-ter truncated KIF5B construct, with the motor domain in N-ter and the dimerization domain in Ct-ter. Dark gray: C-terminal mCherry tag. In red is represented the unstructured N-ter region of kip2 (1-97 a.a) that was added to the N-terminus of truncated kinesin KIF5B (1-560) = Chimera^NKip2^KIF5B. B, C) Kymographs extracted from TIRFm images of kinesin motility assays in presence of KIF5B (50 nM) and ^NKip2^KIF5B chimera (0.5 nM) on GMP-CPP-stabilized MTs spiked with free atto-488 labeled-tubulin (50nM). MT channel (blue), kinesin channel (red) and free-tubulin channel (green). D, E) Quantification of the fluorescence intensity (A.U.) of kinesin accumulation at MT plus-ends (D) and tubulin accumulation at the MT plus-ends (E). F) Quantification of tubulin transport events from the tubulin transport assay for KIF5B and ^NKip2^KIF5B. G) Kymographs extracted from TIRFm images of dynamic MT assays with tubulin at a concentration of 12 µM for the control (Top image) and using the indicated Kip2NKIF5 versions at 10 nM concentration (Bottom image). MT channel (blue), kinesin channel (Red) and tubulin channel (green). H, I) Quantification of MT growth rate (H) and catastrophe frequencies (I) from dynamic MT assays for different concentrations of KIF5B and ^NKip2^KIF5B chimera (mean ± SD of mean). Tubulin concentration: 12µM, Control: only tubulin. J) TIRF microscopy images of curved MT extensions at the plus-ends of dynamic MTs, grown in presence of ^NKip2^KIF5B (10 nM). MT channel (blue), kinesin channel (red) and tubulin channel (green). Pink arrows indicate a curved extension and the white arrow a regular growing MT +end. For (D), (E) and (F) the bar represents the median, the box marks the interquartile range, and the vertical line covers the 95% confidence interval. Statistical test: Mann-Whitney.

Remarkably, ^NKip2^KIF5B was able to bind and transport free tubulin (Fig. 4C, F and Movies 3 and 4), contrary to KIF5B that, apart from occasional motorising aggregates (Fig. 4B, F), did not transport tubulin at comparable motor densities (due to its lower affinity, 10 times higher concentrations of KIF5B were used in the assays in order to achieve comparable motor densities on MTs). In addition, ^NKip2^KIF5B and free tubulin efficiently accumulated at MT plus-ends (Fig. 4C, E and Movie 4), whereas tubulin accumulation at MT ends with KIF5B occurred only sporadically in form of aggregates (Fig. 4B, E and Movie 3).

Finally, unlike its wild type counterpart, the chimeric protein was able to promote MT growth. In dynamic MT assays, ^NKip2^KIF5B decreased the catastrophe frequency and increased the rate of MT polymerization in a concentration-dependent manner (Fig. 4G-I), albeit to lower levels compared to Kip2. In addition, MTs polymerizing in presence of ^NKip2^KIF5B displayed curved protofilament extensions (Fig. 4J), as observed for bKip2-WT (Fig. 3D). In conclusion, the N-terminus of Kip2 is indeed a transferable domain sufficient to promote MT growth when placed in combination with a kinesin motor domain.

## DISCUSSION

Microtubule-associated proteins promote MT growth mainly through a combination of two mechanisms: structural stabilization of MT plus-ends and decrease of the catastrophe rate, as shown for Doublecortin, (Bechstedt *et al*, 2014; Moores *et al*, 2006) or addition of free tubulin at MT plus-ends, exemplified by Stu2/XMAP-215 family (Brouhard *et al*, 2008). Latter proteins act as processive MT polymerases as they remain associated to the MT plus-end during both MT growth and shrinkage, increasing the tubulin polymerization rate, without affecting the rate of catastrophe (Howard & Hyman, 2007). The kinesin Kip2 seems to utilize both mechanisms, since it strongly decreases catastrophes and increases the MT polymerization rate (Hibbel *et al*, 2015; Bowne-Anderson *et al*, 2015).

By performing structure-function analysis, we found that the N-terminal extension preceding the motor domain is required for the MT-growth-promoting activity of Kip2. In vivo, deletion of the 70-terminal amino acids conferred mild sensitivity to benomyl and reduced the length of cMTs (Fig.1F, G). In vitro, the deletion mutant bKip2-Δ70 retained substantial anti-catastrophe activity, but almost completely abolished its capacity to increase the speed of MT polymerization at all concentrations. Thus, the Δ70 deletion largely uncouples the anti-catastrophe from the MT-polymerizing activity of Kip2, suggesting that the Kip2 N-terminus is required to add tubulin to MT plus-ends.

Combining the N-terminal deletion with the P1 mutation in the motor domain completely abolishes the ability of the kinesin to promote MT growth at any concentration (Fig. 1E), implying that next to the N-terminus, the motor domain participates in the MT polymerase activity. In the bKip2-P1 mutant, both anti-catastrophe and polymerase activities are reduced (Fig. 1C, D). However, the fact that bKip2-P1 has lower affinity for the GDP lattice (2,8-fold reduction compared to bKip2-WT, Fig. S1I) complicates drawing conclusions from experiments with dynamic MTs, where the overall MT load of the kinesin is reduced.

Further insight regarding the function of the different domains can been obtained from the behaviour of the different Kip2 versions on GMP-CPP MTs, since all mutants display similar affinity for this lattice (Fig. S1H). Here, bKip2-Δ70 is defective both in kinesin and tubulin accumulation at the plus-end about 3-fold (Fig. 2D, C and Fig. S5D), showing that kinesin and tubulin accumulation at the plus-end are linked and require the Kip2 N-terminus. In contrast, the bKip2-P1 mutant is capable of residing at plus-ends, but is defective in tubulin plus-end accumulation. This suggests that the motor domain participates in retaining free tubulin at the site of its MT incorporation.

Kip2 uses its N-terminus to bind and transport tubulin to MT plus-ends, an observation to our knowledge unprecedented for a mitotic kinesin. Together with the data discussed above, this finding supports a model in which Kip2 promotes MT growth by delivering tubulin to MT plus-ends. We measured tubulin transport events of the order of 1 dimer/minute, at a tubulin concentration of 50nM and Kip2 concentration of 16nM. Using a conservative calculation of a molecular ratio of 1 tubulin dimer:Kip2, at physiological concentrations of 100x more tubulin and 3x more Kip2, tubulin transport could provide almost 20% of the tubulin required for MT growth. Even more tubulin could be transported however, since transport might also occur in the form of droplets (Fig. S3, Movie 2). This type of transport may become relevant in presence of proteins such as Bik1/CLIP-170 and Bim1/EB1 that were shown to form condensates with tubulin (Miesch *et al*, 2023; Meier *et al*, 2023). Therefore, tubulin transport could serve at least as an auxiliary mechanism to promote MT growth.

In further support of this model, the Kip2 motor activity is also required to boost MT polymerization (Fig. 3A-C). Although it still reduces catastrophe, Kip2 in solution and/or bound to MTs in presence of ADP does not increase MT growth speed, indicating that the mobility of the kinesin is required for its polymerase activity. Of note, two major Cdk1 phosphorylation sites that regulate Kip2 activity and MT length *in vivo* are found in the N-terminus (S63) and the motor domain (T275) (Drechsler *et al*, 2015). We therefore think that the N-terminus and the motor domain act in concert to add tubulin to the MT at plus-ends (Fig. 5). The P1 motor site also binds tubulin (Chen *et al*, 2023), but we found that it is not required for tubulin transport along MTs (Fig. 2A, B). Thus, the N-terminus could bind and hand over free tubulin dimers to the motor domain that in turn facilitates their incorporation to a protofilament (Fig. 5). The kinesin may not only add the tubulin that it transports but also capture and add tubulin directly from the solution to the MT end. We cannot think of any obvious reason why one process could be preferred over the other. In agreement with the proposed model, we could frequently observe curved, over-elongated structures at MT plus-ends when MTs polymerized in presence of Kip2, but not when the Kip2 N-terminus was absent (Fig. 3D, E).

**Figure 5:**
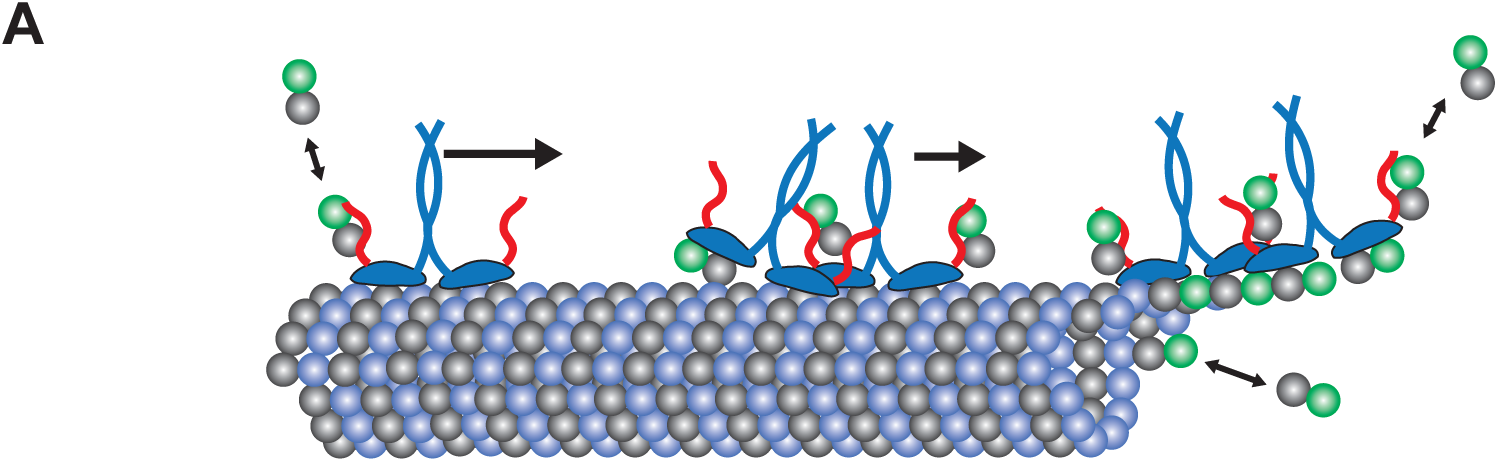
Model of MT polymerization by Kip2. A) Scheme of Kip2 moving alone or in a cluster, capturing and transporting tubulin particles toward the MT plus-end. Kip2 straightening and adding either transported tubulin or directly captured tubulin at the MT plus-end.

Fusion of the Kip2 N-terminus to KIF5B transforms KIF5B to a tubulin-transporting kinesin that promotes MT growth, albeit less efficiently than Kip2. Similar to Kip2, ^NKip2^KIF5B decreases the catastrophe rate and increases the rate of MT growth (Fig. 4G, H, I), parameters that are not affected by KIF5B, that acts rather as a rescue factor (Budaitis *et al*, 2022; Andreu-Carbó *et al*, 2022). This confirms that the Kip2 N-terminus mediates capture of tubulin by the kinesin and shows that the N-terminus is a domain capable of transferring the Kip2 properties to other kinesins, at least partly.

Transport of tubulin by kinesins has been documented since decades (Galbraith & Gallant, 2000) and is required for axon maintenance and regeneration as well as cilia growth. In axons, tubulin synthesized within the cell body is transported in the form of MTs (Hoffman & Lasek, 1980), whereas soluble tubulin is transported to the distal part of the growing axon through a still unknown mechanism. In cilia, kinesin-2 transports tubulin that binds to the intraflagellar transport complex (IFT) to the tip of the cilia (Bhogaraju *et al*, 2013), to promote cilia growth. Here, tubulin transport is considered an auxiliary mechanism, since the major pool of tubulin seems to arrive to the tip of the cilia by diffusion (Craft Van De Weghe *et al*, 2020). In fungi, kinesin-coupled tubulin transport may ensure MT growth and delivery of cellular material and spindle positioning factors to rapidly growing, distal cell ends (Browning *et al*, 2000; Grava & Philippsen, 2010; Estrem *et al*, 2017). Similarly, the kinesin KIF17 couples MT stabilization to plus-end directed transport of the tumour suppressor and spindle positioning factor APC (Jaulin & Kreitzer, 2010), resembling the Kip2-mediated transport of Kar9 and dynein (Maekawa *et al*, 2003; Markus *et al*, 2009; Caudron *et al*, 2008). Interestingly, KIF17 also binds free tubulin with its tail domain, but it is not clear whether it is coupled to transport and MT growth (Acharya *et al*, 2013).

The combination between a kinesin motor domain and a charged, unstructured region that binds tubulin is evident in most Kip2 homologues, even the more distant ones (Lawrence *et al*, 2002; Steinberg, 2007; Raudaskoski, 2022; Wickstead & Gull, 2006). Nevertheless, kinesins bearing motor domains linked to strongly basic, unstructured regions can be often found in many organisms. Combining motility with tubulin-binding and transport seems to be a common module used by MT-growth-promoting kinesins to control MT dynamics. Further studies on the role of tubulin transportation by kinesins are needed to explore whether this mechanism can promote MT growth in environments were diffusion of proteins may be limiting, as for example in the axon of neuronal cells.

## MATERIALS AND METHODS

### Yeast and Bacterial Growth Conditions, Strains and Plasmids

Yeast strains are listed in Supplementary Table 1. Media and genetic manipulations were done as described (Guthrie and Fink, 1991). Standard molecular biology methods were used (Ausubel et al., 1989). Plasmids are listed in Supplementary Table 2. Point mutations (for creation of Kip2P1 mutant) were introduced by site-directed mutagenesis (Phusion polymerase, Thermo Scientific).

### Cloning

Constructs used for yeast transformation were made as follows: the Kip2 coding region, flanked by 379 bp upstream and downstream (379-Kip2-379), was obtained by PCR amplification of the region from genomic *S. cerevisiae* DNA. The amplified sequence was then inserted into the integrative pRS305 vector. The resulting plasmid was pRS305-Kip2. Kip2-Δ70 mutant was derived from the above plasmid.

For kinesin expression in insect cells, the full-length version of Kip2 was obtained by PCR amplification of the region from genomic *S. cerevisiae* DNA. The amplified sequence was inserted into a pFastBac1-mCherry-PrSc-6xHis vector. The resulting plasmid was pFastBac1-Kip2-mCherry-PrSc-6xHis. The Kip2-Δ70 version was derived from the above plasmid.

For kinesin expression in bacteria, the truncated (“bonsai”) version of Kip2 [bKip2-WT (Kip2(1-560))] was obtained by PCR amplification of the region from pFastBac1 vector (see above). The amplified sequence was then inserted into pET-MBP-PrSc-mCherry-6xHis vector, a pET-M43 derivative. The resulting plasmid was pET-MBP-PrSc-Kip2(1-560)-WT-mCherry-6xHis. Kip2-Δ37, Kip2-Δ70, Kip2-P1 and Kip2-P1Δ70 were derived from the above plasmid.

The truncated version of KIF5B [KIF5B(1-560)] from the organism *Mus musculus* was obtained by PCR amplification of the region from the plasmid pKin1B (pKin1B was a gift from Anthony Brown (Uchida *et al*, 2009) Addgene plasmid #31604; http://n2t.net/addgene:31604; <u>RRID:Addgene_31604</u>)). The amplified sequence was inserted into the pET-M43-derived, pET-MBP-PrSc-mCherry-6xHis vector. The resulting plasmid was pET-MBP-PrSc-KIF5B(1-560)-mCherry-6xHis.

pET-MBP-PrSc-Kip2(1-97)-KIF5B(1-560)-mCherry-6xHis (coding for ^NKip2^KIF5B protein) was created by PCR amplification of the Kip2(1-97) region from pET-MBP-PrSc-Kip2(1-560)-mCherry-6xHis and subsequent insertion into pET-MBP-PrSc-KIF5B(1-560)-mCherry-6xHis.

### Incorporation of pS305-Kip2/pRS305-Kip2Δ70 vectors as single copy in yeast cells

Incorporation of the integrative vectors bearing different Kip2 versions (Supplementary Figure S1J) in the yeast genome was examined with primers RS_copy_chk_F/LEU2_copy_chk_1f (annealing at 56°C). Multiple incorporation of the vector in the genome was examined with primers RS_copy_chk_F/RS_copy_chk_2r (annealing at 53°C) and colonies negative for a PCR product were selected. Presence of Kip2/Kip2Δ70 gene was verified with primers 3’ kip2 BamHI/kip2 pro seq f (annealing at 50°C). PCR reactions were done following One*Taq^®^* DNA polymerase protocol. Primer sequences are listed in Supplementary Table 3.

### Live cell imaging

Cells were imaged with Olympus IX83 inverted microscope (Tokyo, Japan) coupled to a Yokogawa W1 spinning disk unit, using an 60X UPLXAPO 1.42NA WD 0.15mm, oil-immersion lens and 488nm laser for sample excitation. Emitted photons were collected by a sCMOS Fusion BT Hamamatsu camera (2304*2304 pixels, 6.5µm pixel size, QEmax 95%). Imaging was performed in a temperature-controlled chamber, at 23°C. z-stacks of 11 slices, with 0.5µm step/slice, from 7 positions were collected every 70s, for a total time of 12 min (10 cycles), using the CellSens software.

### Protein purification

Tubulin was isolated from pig brain by two cycles of microtubules polymerization/depolymerization in high molarity pipes buffer as described in Castoldi and Popov (2003).

MBP-PrSc-Kip2(1-560)-WT-mCherry-PrSc-His and the Δ37, Δ70, P1, P1Δ70 and MBP-KIF5B(1-560)-mCherry-6xHis and MBP-Kip2(1-97)-KIF5B(1-560)-mCherry-6xHis constructs were expressed in *E. coli* (Rosetta strain). Bacteria were grown in TB medium at 37°C until culture reached OD 0.6-0.7 and recombinant protein expression was triggered by the addition of 200 µM IPTG for 16 h at 16°C. Cells were harvested, resuspended in purification buffer for MBP-Kip2-mCherry-His (2XH buffer: 40mM HEPES, 400mM KCl, 2mM MgCl_2_, 1mM ATP, pH 7.4) with 5mM β-mercaptoethanol and protease inhibitors (SigmaFAST, protease inhibitor Cocktail EDTA-Free, S8830) and lysed by sonication. The proteins were first bound to Ni-NTA resin, then eluted with 300mM Imidazole and then bound to an Amylose resin (E8021; NEB) and eluted with 10mM maltose, followed by cleavage with PreScission protease overnight to remove the MBP-tag. The free MBP-tag was removed using amylose resin and the GST-PreScission was removed using a Glutathione Sepharose 4B resin. The proteins were further purified by size exclusion chromatography using the Akta Start system with a HiPrep™ 16/60 Sephacryl® S-300 HR column (Cytiva). The proteins were aliquoted, flash frozen in liquid nitrogen, and stored at −80°C.

### Dynamic instability assay

Tubulin was labelled with NHS-Biotin (Ez-link NHS-LC-LC-Biotin, 21343; Thermo Fisher Scientific), NHS-ATTO-488 (AD488-31; ATTO-TEC), NHS-ATTO-647 (AD647-31; ATTO-TEC) during a cycle of polymerization/depolymerization as described in (Hyman *et al*, 1991). MT seeds were polymerized from 20µM tubulin (with 5% of tubulin-ATTO-647 and 5% tubulin-biotin) with 1mM Guanosine-5’-[(α,β)-methyleno]triphosphate (GMPCPP) in PEM80 (80 mM PIPES, 1 mM EGTA, 1 mM MgCl_2_, pH 6.9), at 37°C for 30 min, centrifuged at 26,000 g for 15 min and resuspended in PEM80. Glass coverslips were cleaned and silanized as described in (Brouhard *et al*, 2008). The flow cells were assembled from the silanized coverslip and a coverglass using double-sided tape (LIMA, 70pc). The flow cell was treated with 50 µg/ml of neutravidin in PEM80 and then washed with a solution of 2% Pluronic F-127. The flow cell was then washed with PEM80 and MT seeds were allowed to attach to the neutravidin-coated surface. Next, for Kip2-WT and Kip2-Δ70, the polymerization mix containing 8µM tubulin (supplemented with 5% tubulin-ATTO-488) and 1mM GTP in PEM20 buffer (20mM PIPES, 1mM EGTA, 1mM MgCl_2_, pH 6.9) supplemented with 100mM KCl, 0.01% Tween-20, 0.05% methyl-cellulose, 20mM glucose, 20µg/ml glucose oxidase, 8µg/ml catalase, 0.1mg/ml BSA, 1mM DTT, 1mM GTP, and 1mM ATP, with 5nM of Kip2-WT full-length or Kip2-Δ70 was flowed in. For bKip2 constructs the polymerization mix containing 8µM tubulin (supplemented with 5% tubulin-ATTO-488) and 1mM GTP in HEM100 buffer (100mM HEPES, 100mM KCl, 10mM MgCl_2_, 1mM EGTA, pH 6.9) supplemented with 100mM KCl, 0.01% Tween-20, 0.05% methyl-cellulose, 20mM glucose, 20µg/ml glucose oxidase, 8µg/ml catalase, 0.1mg/ml BSA, 1mM DTT, 1mM GTP, and 1mM ATP was flowed in. Increasing concentrations of bKip2-WT or the different constructs were added to test the concentration-dependent effect of the kinesins on MT polymerization *in vitro*. The MT polymerization in different conditions was recorded for 10 min, at 30°C, using an objective-based azimuthal ilas2 TIRF microscope, Nikon Eclipse Ti, modified by Roper Scientific and an Evolve 512 camera from Photometrics, driven by Metamorph software. Time-lapse recording was performed every 5s, for 10 min for the 488, 565 and 647nm laser lines.

For ^NKip2^KIF5B dynamic instability assays, chambers were washed with PEM80 and then the polymerization mix containing 12µM free tubulin (with 5% ATTO-488 tubulin) in PEM60 (60mM PIPES, 1mM MgCl_2_, 1mM EGTA pH 6.9), supplemented with 0.01% Tween-20, 0.05% methyl-cellulose, 2mg/ml BSA, 20mM DTT, 4.5mg/ml glucose, 0.2mg/ml glucose oxidase, 0.35mg/ml catalase, 2mM ATP, 2mM GTP and supplemented with 3, 5, 10 or 20nM ^NKip2^KIF5B. The chamber was fully-sealed and immediately imaged by TIRF in a temperature-controlled chamber at 30°C. Data was collected every 3s, for 10min.

### Kip2 motility assay

GMPCPP-stabilized MTs were polymerized from 20µM tubulin (with 5% of tubulin-ATTO-488 and 5% tubulin-biotin) with 0.5mM GMPCPP in PEM80, at 37°C for 1h. Then the GMPCPP-stabilized MTs were attached to the flow chamber coverslip functionalized with neutravidin and passivated with pluronic-F-127 as described above. 0.1nM of Kip2-WT or Kip2-Δ70 in PEM20 buffer (20mM PIPES, 1mM MgCl_2_, 1mM EGTA, pH 6.9), supplemented with 100mM KCl, 0.01% Tween-20, 20mM glucose, 20µg/ml glucose oxidase, 8µg/ml catalase, 0.1mg/ml BSA, 1mM DTT, 1mM GTP, and 1mM ATP was flowed in. Alternatively, 0.5 to 5nM of bKip2-WT, bKip2-Δ37, bKip2-Δ70, bKip2-P1 and bKip2-P1Δ70 in HEM100 buffer (100mM HEPES, 10mM MgCl_2_, 1mM EGTA, pH 6.9), supplemented with 100mM KCl, 0.01% Tween-20, 20mM glucose, 20µg/ml glucose oxidase, 8µg/ml catalase, 0.1mg/ml BSA, 1mM DTT, 1mM GTP, and 1mM ATP, was flowed in the flow chamber. Time-lapse recording every 5s or 2s, for 10 min, at 642 and 565nm was performed using Metamorph software.

For KIF5B and ^NKip2^KIF5B TIRF microscopy motility assay, the same procedures as above were followed. Long microtubules instead of microtubule seeds were used. Motility buffer: PEM20 (20mM PIPES, 1mM MgCl_2_, 1mM EGTA, pH 6.9) supplemented with 0.01% Tween-20, 0.05% Methyl cellulose, 2mg/ml BSA, 20mM DTT, 4.5mg/ml glucose, 0.2mg/ml glucose oxidase, 0.35mg/ml catalase and 2mM ATP, supplemented with 20nM KIF5B or 0.5nM ^NKip2^KIF5B was added. Imaging was performed in a temperature-controlled chamber, at 30°C. Data was collected every 2s, for 8 min, using Metamorph software

### Kip2 affinity assay

GMPCPP-stabilized MTs were polymerized from 20µM tubulin (with 5% of tubulin-ATTO-488 and 5% tubulin-biotin) with 0.5mM GMPCPP in PEM80, at 37°C, for 1h. MTs stabilized with Taxotere were polymerized from 20µM tubulin (with 5% of tubulin-ATTO-488 and 5% tubulin-biotin) with 1mM GTP in PEM80, at 37°C, for 45 min. Then, 1µM Taxotere was added, and an additional 5µM Taxotere was re-added every 15 min, at 37°C to a final Taxotere concentration of 20µM. GMPCPP-stabilized or Taxotere-stabilized MTs were attached to the flow chamber coverslip functionalized with neutravidin and passivated with pluronic-F-127 as described above. Then, 0.1 to 10nM Kip2-WT or Kip2-Δ70, in PEM20 buffer (20mM PIPES, 1mM MgCl2, 1mM EGTA, pH 6.9), supplemented with 100mM KCl, 0.01% Tween-20, 20mM glucose, 20µg/ml glucose oxidase, 8µg/ml catalase, 0.1mg/ml BSA, 1mM DTT, and 1mM ATP were flowed. Alternatively, 0.5 to 50nM of bKip2-WT, bKip2-Δ70, bKip2-P1 and bKip2-P1Δ70 in HEM100 buffer (100mM HEPES, 10mM MgCl_2_, 1mM EGTA, pH 6.9), supplemented with 100mM KCl, 0.01% Tween-20, 20mM glucose, 20µg/ml glucose oxidase, 8µg/ml catalase, 0.1mg/ml BSA, 1mM DTT, 1mM GTP, and 1mM ATP was flowed in the flow chamber. Images for MTs and Kip2 were recorded at 642 and 565nm, respectively.

### Tubulin binding and transport assay

For Kip2-WT: GMPCPP-stabilized MTs were polymerized from 20µM tubulin (with 5% of tubulin-ATTO-488 and 5% tubulin-biotin) with 0.5mM GMPCPP in PEM80, at 37°C, for 1 h. Then the GMPCPP-stabilized MTs were attached to the surface of a flow chamber functionalized with neutravidin and passivated with pluronic-F-127 as described above. Then, 10nM of Kip2-WT was incubated 5 min on ice with 1mM AMP-PNP in PEM20 buffer (20mM PIPES, 1mM MgCl_2_, 1mM EGTA, pH 6.9), supplemented with 100mM KCl, 0.01% Tween-20, 0.1mg/ml BSA, 1mM DTT, and then flowed onto the attached MTs and incubated at 30°C, for 2 min. Then the chamber was washed with PEM20 buffer supplemented with 100mM KCl, 0.01% Tween-20, 0.1mg/ml BSA, 1mM DTT and 1mM ATP. Then, 1µM of ATTO-488 labelled tubulin in PEM80 was flowed in the chamber and incubated for 1 min, then washed away with PEM80 buffer. The motility buffer was subsequently flowed in the chamber to trigger Kip2 motility: PEM20 buffer (20mM PIPES, 1mM MgCl_2_, 1mM EGTA, pH 6.9), supplemented with 100mM KCl, 0.01% Tween-20, 20mM glucose, 20µg/ml glucose oxidase, 8µg/ml catalase, 0.1mg/ml BSA, 1mM DTT, and 10mM ATP. Kinesin movement and tubulin attachment and transport were immediately imaged in TIRF microscopy. After a few minutes Kip2 exchanged AMP-PNP with ATP and began to move on MTs.

For mixed GMPCPP and GDP lattice, MTs were polymerized from 20µM of tubulin with Taxotere and then elongated with 0.5mM GMPCPP (with 5% of tubulin-ATTO-488 and 5% tubulin-biotin), in PEM80, at 37°C, for 1 h. Then, the mixed GMPCPP/Taxotere-stabilized MTs were attached to the surface of a flow chamber functionalized with neutravidin and passivated with pluronic-F-127 as described above. Then, 4 to 20nM of bKip2-WT, bKip2-Δ70 and bKip2-P1 in PEM20 buffer (20mM PIPES, 1mM MgCl_2_, 1mM EGTA, pH 6.9) or HEM100 buffer (100mM HEPES, 1mM MgCl_2_, 1 mM EGTA, pH 6.9) supplemented with 50 or 100mM KCl, 0.01% Tween-20, 20mM glucose, 20µg/ml glucose oxidase, 8µg/ml catalase, 0.1mg/ml BSA, 1mM DTT, 1mM GTP or GDP, 1mM ATP, and 50nM of free ATTO-488 labelled tubulin was flowed in the flow chamber. Time-lapse recording every 5s or 2s, for 10 min at 488, 565 and 642nm was performed using Metamorph software.

For KIF5B and ^NKip2^KIF5B tubulin transport assays, long microtubules instead of microtubule seeds were used. Tubulin transport buffer: PEM20 (20mM PIPES, 1mM MgCl_2_, 1mM EGTA, pH 6.9), supplemented with 0.01% Tween-20, 0.05% Methyl cellulose, 2mg/ml BSA, 20mM DTT, 4.5mg/ml glucose, 0.2mg/ml glucose oxidase, 0.35mg/ml catalase, 2mM ATP, 2mM GTP and 50nM ATTO-488 tubulin, supplemented with 50nM KIF5B or 5nM ^NKip2^KIF5B. Imaging was performed in a temperature-controlled chamber, at 30°C. Data was collected every 2s, for 10 min.

### Electron microscopy

MTs stabilized with GMPCPP were polymerized with 20µM tubulin with 0.5mM GMPCPP in PEM80, at 37°C, for 1 h. Control MT alone were diluted in PEM80 buffer to a final concentration of 1µM. Kip2-WT was incubated with the GMPCPP-stabilized MTs at a 1:2 ratio for MT:Kip2, in PEM20 buffer (20mM PIPES, 1mM MgCl2, 1mM EGTA, pH 6.9), supplemented with 100mM KCl, 1mM DTT and 1mM ATP or 1mM AMP-PNP.

Negative staining: Samples were dropped off on carbon-coated formvar copper grids (FCF2010-CU, EMS, Hatflied, PA, USA) for 2 min. The excess liquid was then removed and the grids stained with uranyl acetate 1% for 1 min. The specimens were observed on a Tecnai G2 F20 (200 kV FEG) TEM.

### Image and data analysis

All image analysis was performed using Fiji. Image stacks were corrected for drift using the plugin Fast4Dreg. Dynamic instability parameters were extracted from movies using kymographs. Kymographs were obtained using the plugin Multiple Kymographs and velocity of kinesins or tubulin transports were measured using the macro velocity measurement (developed by Volker Bäcker https://dev.mri.cnrs.fr/projects/imagej-macros/wiki/Kymograph). For fluorescence intensity quantification of Kip2 and tubulin along the lattice, a polyline (width of 3 pixels) was drawn on the MT and the initial fluorescence was measured at the first frame. For the quantification of accumulation of kinesin or tubulin at the MT plus-end, a circle (diameter of 6 pixels) was drawn at the MT plus-end (MT plus-end was generally indicated by the plus-end accumulation of the kinesin).

### Statistical analysis

All data were processed using Excel or GraphPad Prism 5 software. The number of experiments and/or samples are indicated in the figure legend. The SD is shown either in the graphs, or as indicated. N.S. (not significant) or asterisks indicate P values. Mann-Whitney or unpaired t-test were used for the calculation of statistical significance (refer to Excel sheet of supplementary data).

## AUTHOR CONTRIBUTIONS

S.N., D.P., D.L. designed the experiments. S.N., D.P carried out the experiments. A.N. did the electron microscopy imaging. S.N., D.P., A.A., D.L. analyzed the data. D.P., D.L. developed the model. H.D., D.P., D.L. wrote the manuscript. D.P., D.L. initiated the research and supervised the work. All authors discussed the results and commented on the manuscript.

## ACKNOWLEDGEMENTS

We acknowledge the imaging facility MRI, member of the national infrastructure France-BioImaging infrastructure supported by the French National Research Agency (ANR-10-INBS-04, «Investments for the future». D. Liakopoulos acknowledges the support of the French Agence Nationale de la Recherche, grant ANR-14-CE09-0014-01 (ReconstMT-Act) and D. Portran acknowledges the support of ARC, grant ARF20161236770. S. Nadalis acknowledges the support of the A. G. Leventis Foundation and the CBS2 Doctoral School of the University of Montpellier.

## Figure and Movie Legends

For details on statistics See also *Supplementary Statistical Table 1*. N refers to the number of independent experiments.

**Figure S1.**
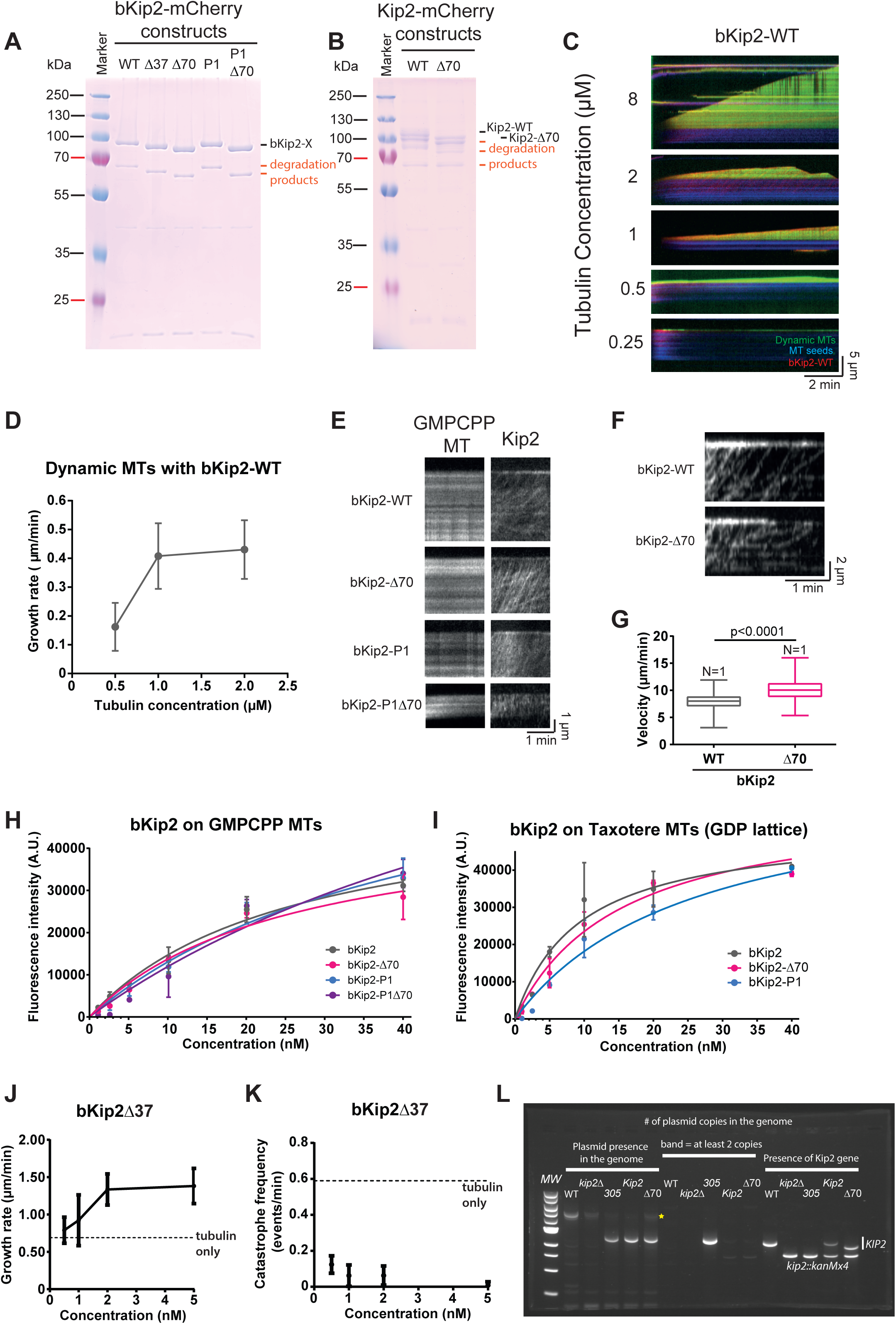
A) , B) Coomassie stained gels of bacterially purified bKip2 and full-length Kip2-WT versions purified from SF9 cells. C) Kymographs extracted from TIRFm images of dynamic MT assays in presence of 1nM of bKip2-WT and varying tubulin concentrations at the sub-polymerization regime (0.25-2µM). MT growth takes place already at 0.5µM tubulin. Blue: MT seeds, red: kinesin and green: atto-488-tubuli. D) Quantification of MT growth rate as a function of tubulin concentration shown in (C) (mean ± SD of mean). E) Kymographs extracted from motility assays of bKip2 mutants (5nM) on GMP-CPP-stabilized MTs. F) Kymographs extracted from motility assays of bKip2 and bKip2-Δ70 (only the kinesin channel is shown). G) Quantification of velocity from motility assays of bKip2 and bKip2-Δ70. The bar represents the median, the box marks the interquartile range, and the vertical line covers the 95% confidence interval. Statistical test: Mann-Whitney. H, I) Quantification of kinesin fluorescence (A.U.) on MTs on GMPCPP-stabilized MTs (H) and Taxotere-stabilized MTs (I) (mean ± SD of mean). Curves for all bKip2 mutants have been fitted assuming one-site specific binding and no cooperativity. For bKip2-WT, bKip2-Δ70 and bKip2-P1 at least 50 MTs were analyzed for each concentration, and experiments were repeated 3 times. J) MT growth rate as a function of bKip2-Δ37 concentration in presence of 8 µM tubulin (mean ± SD of mean). The dashed line shows the MT growth rate at 8 µM tubulin concentration in absence of kinesin. A single experiment for each concentration was performed. K) MT catastrophe as a function of bKip2-Δ37 concentration in presence of 8 µM tubulin (mean ± SD of mean). The dashed line shows the MT catastrophe frequency at 8 µM tubulin in absence of kinesin. A single experiment for each concentration was performed. L) PCR to test the number of integrated plasmid copies in the yeast mutants used for the measurement of the cMT lengths in vivo in Fig. 1G.

**Figure S2.**
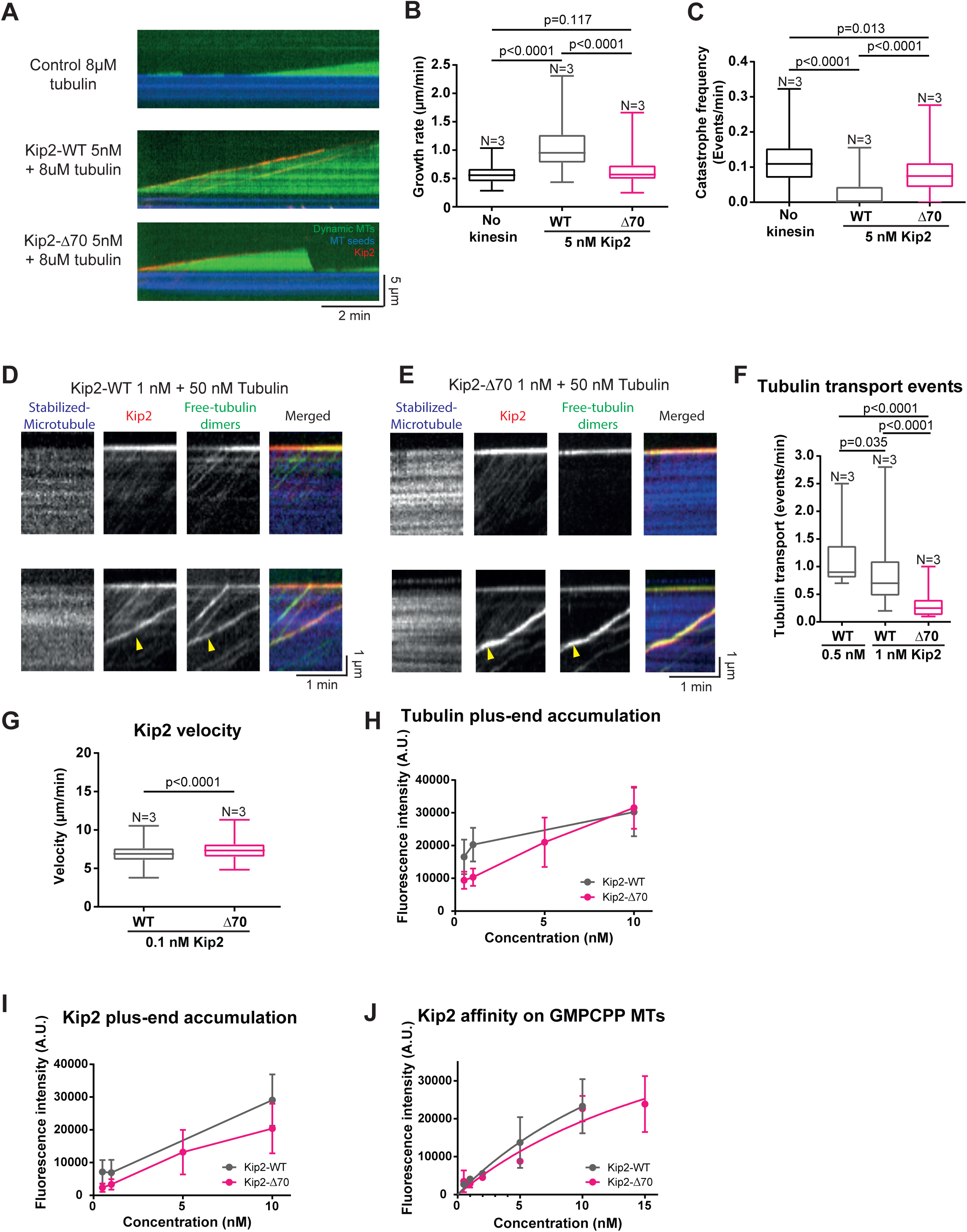
A) Kymographs extracted from dynamic MT assays using the full length Kip2-WT and Kip2-Δ70-versions at 5nM concentration. Blue: MT seeds, red: kinesin and green: atto-488-tubulin. B, C) Quantification of catastrophe frequency and MT growth rate in presence of full length Kip2-WT and Kip2-Δ70 at 5nM concentration. Tubulin concentration in the assay: 8 µM. Control: tubulin only. D, E) Kymographs extracted from motility assays of full length Kip2-WT and Kip2-Δ70 (both at 5nM), on stabilized MTs spiked with free 50nM of atto-488 labeled-tubulin. Blue: MT seeds, red: kinesin and green: atto-488-tubulin. Note the green tubulin traces that colocalize with kinesin, only large clusters are transported by full-length Kip2-Δ70. F) Quantification of tubulin transportation events per min, per MT, for Kip2-WT) in the assays in (D) and (E). G) Median velocity of Kip2-WT and Kip2-Δ70 particles at 0.1 nM. H, I) Quantifications of tubulin fluorescence intensity (A.U.) and Kip2 fluorescence intensity (A.U.) at the MT plus-end for Kip2-WT and Kip2-Δ70 at concentrations of 1, 2, 5 and 10 nM (at least 139 MTs were analyzed per conditions and the experiment was repeated at least 3 times) (mean ± SD of mean). J) Quantification of kinesin fluorescence (A.U.) on GMPCPP-stabilized MTs at 1, 2, 5 and 10 nM of Kip2. Curves have been fitted assuming one-site specific binding and no cooperativity. At least 200 MTs were analyzed for each concentration, and experiments were repeated 3 times (mean ± SD of mean). For (B), (C), (F) and (G) the bar represents the median, the box marks the interquartile range, and the vertical line covers the 95% confidence interval. Statistical test: Mann-Whitney.

***Figure S3***

A) Sequence images of full length Kip2-WT (10nM) attached in presence of AMPPNP on GMPCPP-stabilized MTs, subsequently incubated with free atto-488 labeled-tubulin (1 µM), followed by kinesin ‘release’ with ATP and GTP. Arrows show droplet-like structures (arrows) that merge and move to the tips of MTs. Blue: MT seeds, red: kinesin and green: atto-488-tubulin. Scale bar = 5µm.

B) Kymographs extracted from (A) double headed arrow lines indicated with AMP-PNP and ATP depict the times of immobilization and kinesin release, respectively.

**Figure S4.**
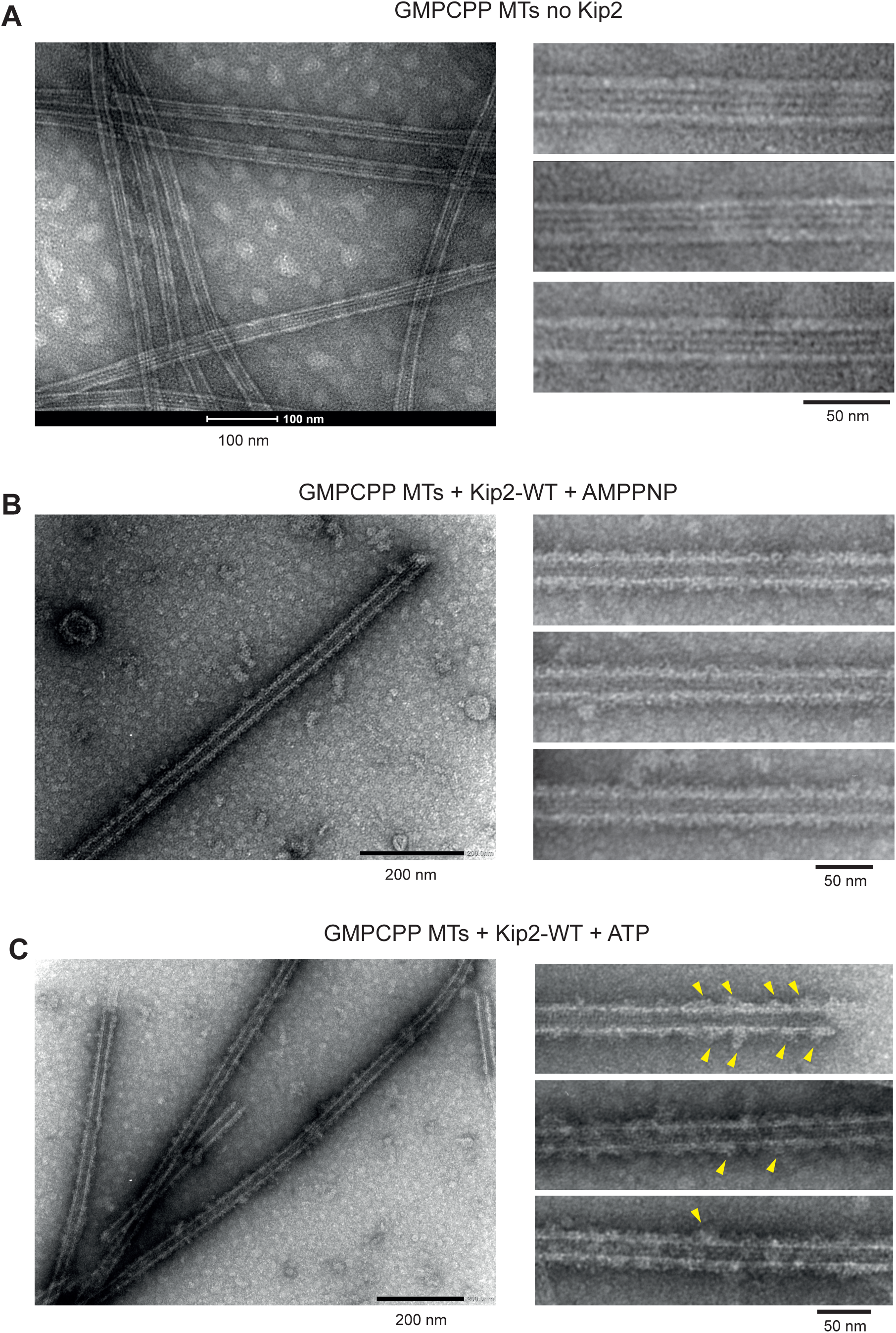
A) Electron microscopy image of a negative-staining control microtubule sample without kinesin, left large field image and right zoom section of microtubules lattices. Left image scale bar: 100 nm. B) Electron microscopy image of a negative-staining microtubule sample in presence of Kip2-WT with AMP-PNP, left large field image and right zoom section of microtubules lattices. Left image scale bar: 200 nm. C) Electron microscopy image of a negative-staining microtubule sample in presence of Kip2-WT with ATP, left large field image and right zoom section of microtubules lattices. Arrows show clusters/accumulation of Kip2 on the MT lattice. Left image scale bar: 200 nm.

**Figure S5.**
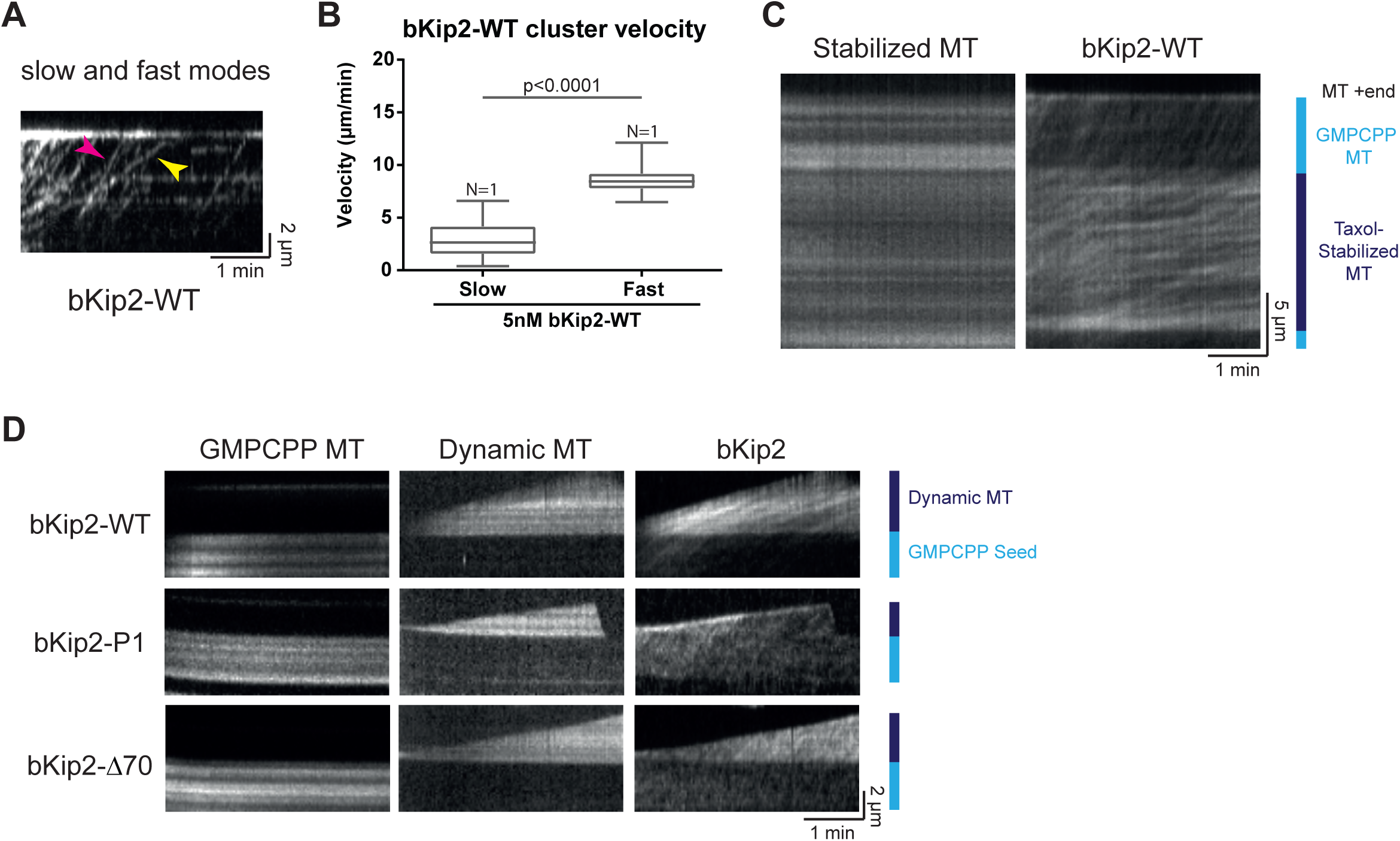
A) Kymograph extracted from TIRFm images of motility assays for bKip2-WT (5 nM) on GMP-CPP-stabilized MTs. Fast mode indicated by pink arrow and slow clustered mode indicated by yellow arrow. Only the kinesin channel is shown. B) Quantification of bKip2-WT velocity for fast and slow particles (for 5nM, NMTs= 25, Nxp= 1). The bar represents the median, the box marks the interquartile range, and the vertical line covers the 95% confidence interval. Statistical test: Mann-Whitney. C) Kymographs extracted from TIRFm images of different bKip2 constructs on mixed lattice GMP-CPP/taxotere-stabilized MTs showing differences of affinity and velocity in dependence of the lattice. Scheme on the right is showing the limit of the GMPCPP *vs* GDP lattice. D) Kymographs extracted from TIRFm images of bKip2-WT, -P1 and -Δ70 motility on dynamic MTs showing differences of affinity and velocity in dependence of the lattice (GMP-CPP for the MT seed and GDP lattice for the dynamic MT). Scheme on the right is showing the limit of the GMP-CPP *vs* GDP lattice.

**Figure S6.**
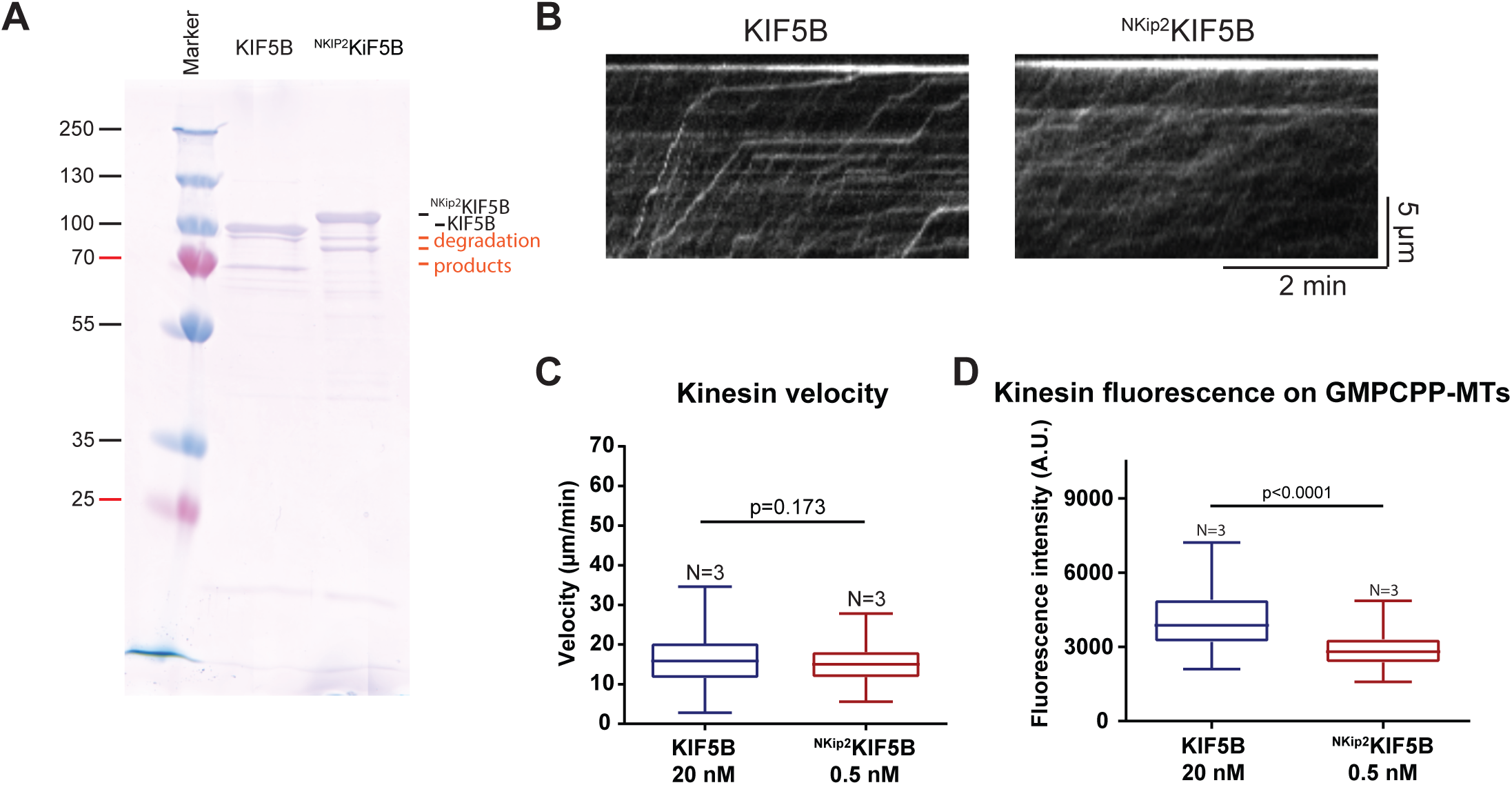
A) Coomassie stained gels of the purified KIF5B and chimeric construct ^NKip2^KIF5B. B) Kymographs extracted from TIRFm images of KIF5B (20 nM) (Left) and ^NKip2^KIF5B (0.5 nM) (Right) motility on GMP-CPP-stabilized MTs. Only the kinesin channel is shown. C) Quantification of KIF5B and ^NKip2^KIF5B chimera velocity from kinesin motility TIRFm assays. D) Quantification of kinesin fluorescence on stabilized MTs from the assays in figure 4A. For (C) and (D), the bar represents the median, the box marks the interquartile range, and the vertical line covers the 95% confidence interval. Statistical test: Mann-Whitney.

***Supplementary Movie 1***

Movie extracted from TIRFm images from tubulin transport assays of bKip2 mutants (16 nM) on stabilized MTs spiked with free atto-488 labeled-tubulin (50nM). MT channel (blue), kinesin channel (red) and free-tubulin channel (green). Scale bar = 5 µm.

***Supplementary Movie 2***

Movie extracted from TIRFm images from tubulin transport assays of Kip2 full-length (10 nM) bound with AMPPNP on GMP-CPP stabilized MTs, then washed and incubated with with free atto-488 labeled-tubulin (1µM) then washed with a buffer containing 10 mM ATP to start the kinesin movement. The movie starts after the last wash with ATP. MT channel (blue), kinesin channel (red) and free-tubulin channel (green). Scale bar = 5 µm.

***Supplementary Movie 3***

Movie extracted from TIRFm images from tubulin transport assays of bKIF5B (50 nM) on stabilized MTs (GMP-CPP-stabilized MTs) spiked with with free atto-488 labeled-tubulin (50nM). kinesin channel (Left panel) and free-tubulin channel (Right panel). Scale bar = 5 µm.

***Supplementary Movie 4***

Movie extracted from TIRFm images from tubulin transport assays of bonsaï ^NKip2^KIF5B (5 nM) on stabilized MTs (GMP-CPP-stabilized MTs) spiked with with free atto-488 labeled-tubulin (50nM). Kinesin channel (Left panel) and free-tubulin channel (Right panel). Scale bar = 5 µm.

***Supplementary Movie 5***

Movie extracted from TIRFm images from MT dynamic assay with 10 nM bonsaï ^NKip2^KIF5B, with GMP-CPP-stabilized seeds in blue, Kinesin in red and polymerizing MTs in green. Scale bar = 5 µm.

---

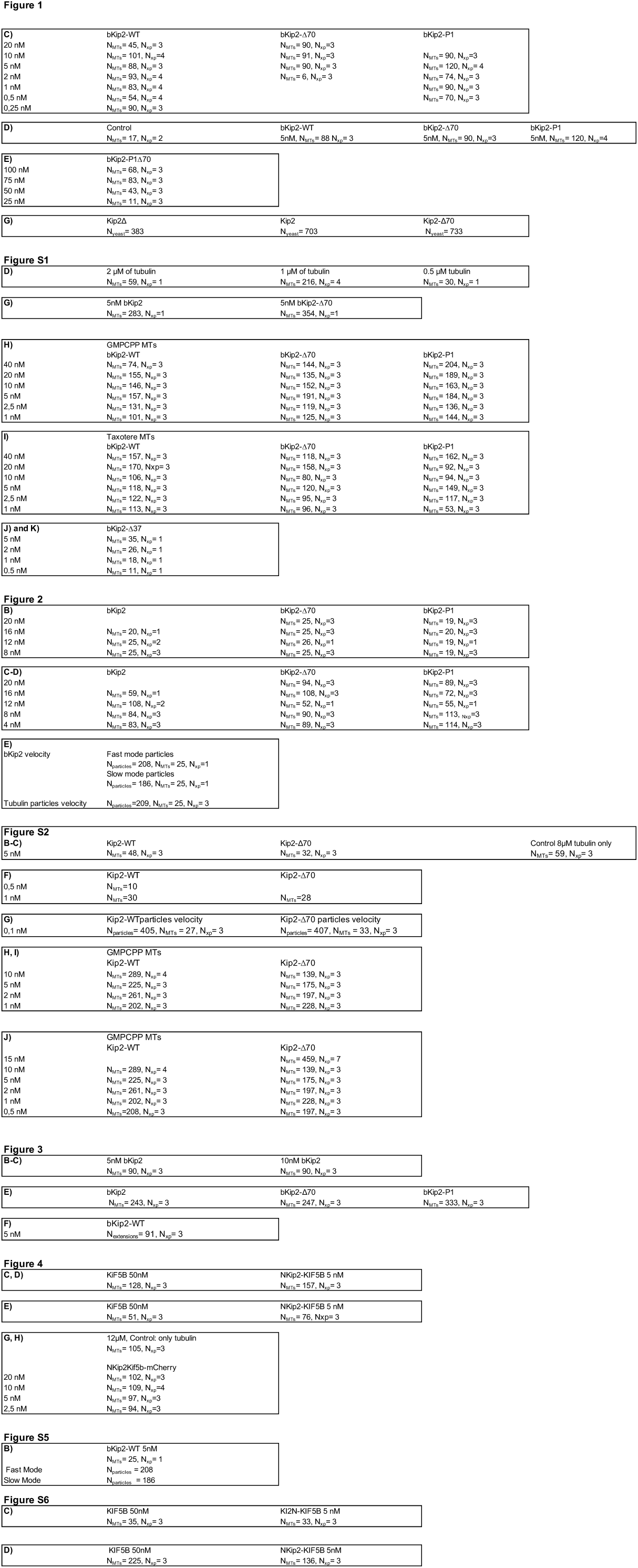

**Table S1.**
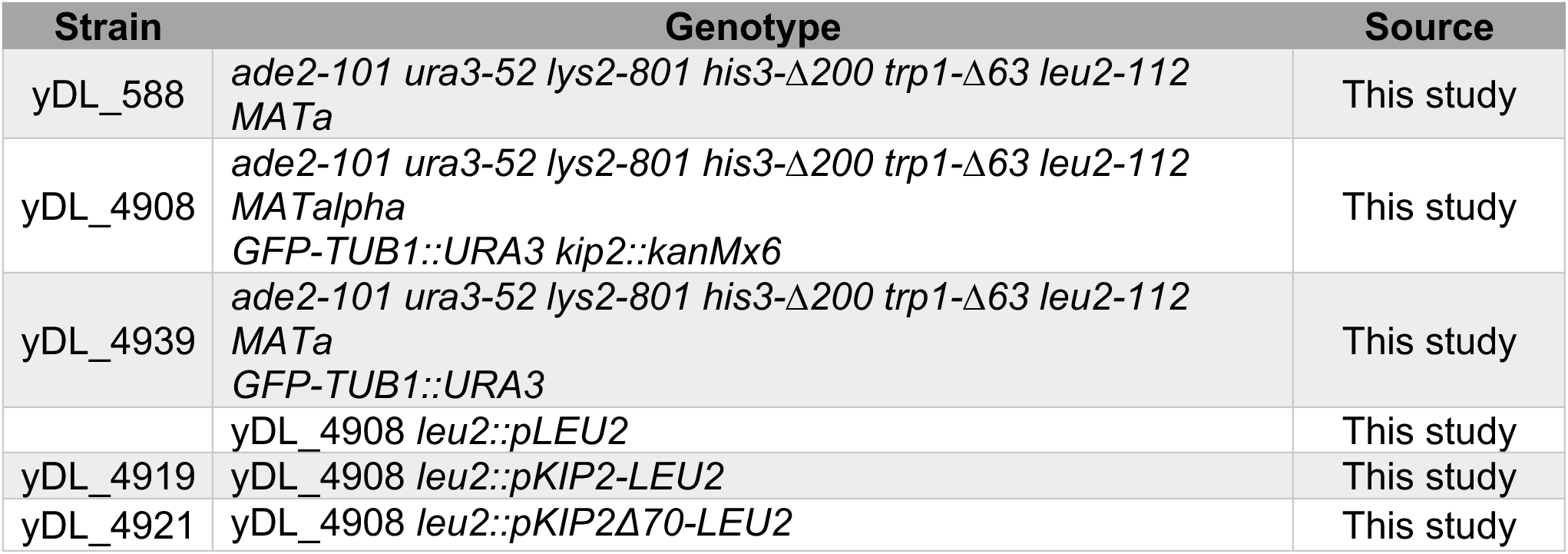
Strains.

| Strain | Genotype | Source |
| --- | --- | --- |
| yDL_588 | *ade2-101 ura3-52 lys2-801 his3-Δ200 trp1-Δ63 leu2-112 MATa* | This study |
| yDL_4908 | *ade2-101 ura3-52 lys2-801 his3-Δ200 trp1-Δ63 leu2-112 MATalpha GFP-TUB1::URA3 kip2::kanMx6* | This study |
| yDL_4939 | *ade2-101 ura3-52 lys2-801 his3-Δ200 trp1-Δ63 leu2-112 MATa GFP-TUB1::URA3* | This study |
|  | yDL_4908 *leu2::pLEU2* | This study |
| yDL_4919 | yDL_4908 *leu2::pKIP2-LEU2* | This study |
| yDL_4921 | yDL_4908 *leu2::pKIP2Δ70-LEU2* | This study |

**Table S2.**
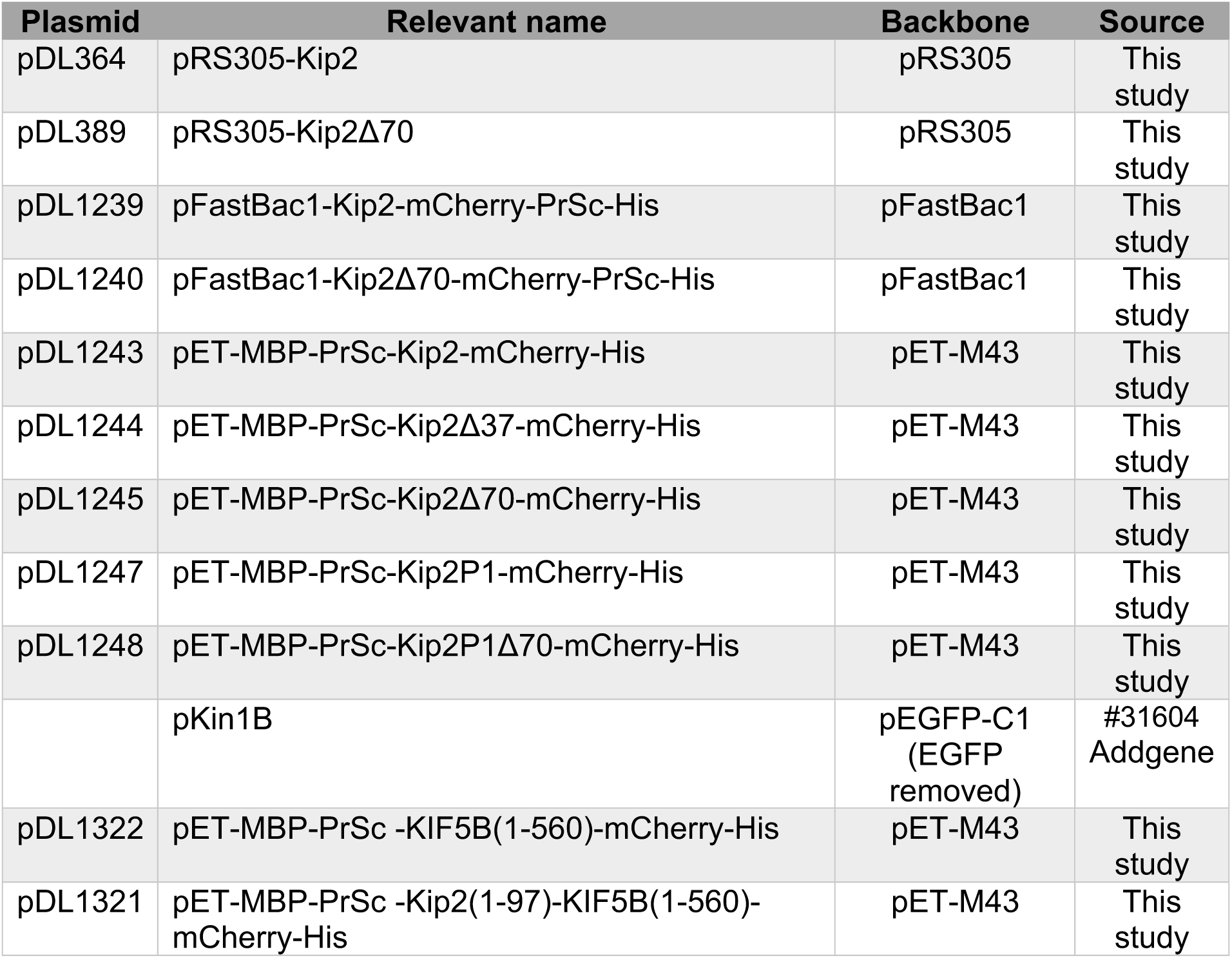
Plasmids.

| Plasmid | Relevant name | Backbone | Source |
| --- | --- | --- | --- |
| pDL364 | pRS305-Kip2 | pRS305 | This study |
| pDL389 | pRS305-Kip2Δ70 | pRS305 | This study |
| pDL1239 | pFastBac1-Kip2-mCherry-PrSc-His | pFastBac1 | This study |
| pDL1240 | pFastBac1-Kip2Δ70-mCherry-PrSc-His | pFastBac1 | This study |
| pDL1243 | pET-MBP-PrSc-Kip2-mCherry-His | pET-M43 | This study |
| pDL1244 | pET-MBP-PrSc-Kip2Δ37-mCherry-His | pET-M43 | This study |
| pDL1245 | pET-MBP-PrSc-Kip2Δ70-mCherry-His | pET-M43 | This study |
| pDL1247 | pET-MBP-PrSc-Kip2P1-mCherry-His | pET-M43 | This study |
| pDL1248 | pET-MBP-PrSc-Kip2P1Δ70-mCherry-His | pET-M43 | This study |
|  | pKin1B | pEGFP-C1<br>(EGFP removed) | #31604<br>Addgene |
| pDL1322 | pET-MBP-PrSc -KIF5B(1-560)-mCherry-His | pET-M43 | This study |
| pDL1321 | pET-MBP-PrSc -Kip2(1-97)-KIF5B(1-560)-mCherry-His | pET-M43 | This study |

**Table S3.**
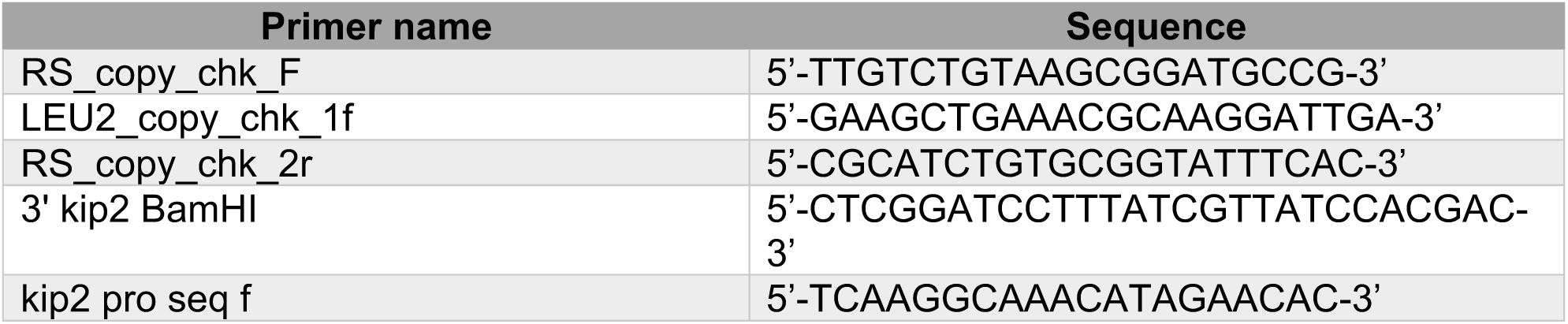
Primers.

## REFERENCES

1. Acharya BR, Espenel C & Kreitzer G (2013) Direct Regulation of Microtubule Dynamics by KIF17 Motor and Tail Domains. J Biol Chem 288: 32302–32313

2. Andreu-Carbó M, Fernandes S, Velluz MC, Kruse K & Aumeier C (2022) Motor usage imprints microtubule stability along the shaft. Dev Cell 57: 5–18.e8

3. Bechstedt S, Lu K & Brouhard GJ (2014) Doublecortin Recognizes the Longitudinal Curvature of the Microtubule End and Lattice. Curr Biol 24: 2366–2375

4. Bhogaraju S, Cajanek L, Fort C, Blisnick T, Weber K, Taschner M, Mizuno N, Lamla S, Bastin P, Nigg EA, et al (2013) Molecular Basis of Tubulin Transport Within the Cilium by IFT74 and IFT81. Science (80- ) 341: 1009–1012

5. Bowne-Anderson H, Hibbel A & Howard J (2015) Regulation of Microtubule Growth and Catastrophe : Unifying Theory and Experiment. Trends Cell Biol 25: 769–779

6. Brouhard GJ, Stear JH, Noetzel TL, Al-Bassam J, Kinoshita K, Harrison SC, Howard J & Hyman AA (2008) XMAP215 Is a Processive Microtubule Polymerase. Cell 132: 79–88

7. Browning H, Hayles J, Mata J, Aveline L, Nurse P & McIntosh JR (2000) Tea2p is a kinesin-like protein required to generate polarized growth in fission yeast. J Cell Biol 151: 15–28

8. Budaitis BG, Badieyan S, Yue Y, Blasius TL, Reinemann DN, Lang MJ, Cianfrocco MA & Verhey KJ (2022) A kinesin-1 variant reveals motor-induced microtubule damage in cells. Curr Biol 32: 2416–2429.e6

9. Carvalho P, Gupta ML, Hoyt MA & Pellman D (2004) Cell cycle control of kinesin-mediated transport of Bik1 (CLIP-170) regulates microtubule stability and dynein activation. Dev Cell 6: 815–29

10. Caudron F, Andrieux A, Job D & Boscheron C (2008) A new role for kinesin-directed transport of Bik1p (CLIP-170) in Saccharomyces cerevisiae. J Cell Sci 121: 1506–13

11. Chen G-Y, Cleary JM, Asenjo AB, Chen Y, Mascaro JA, Arginteanu DFJ, Sosa H & Hancock WO (2019) Kinesin-5 Promotes Microtubule Nucleation and Assembly by Stabilizing a Lattice-Competent Conformation of Tubulin. Curr Biol 29: 2259–2269.e4

12. Chen X, Portran D, Widmer LA, Stangier MM, Czub MP, Liakopoulos D, Stelling J, Steinmetz MO & Barral Y (2023) The motor domain of the kinesin Kip2 promotes microtubule polymerization at microtubule tips. J Cell Biol 222: e202110126

13. Cottingham FR & Hoyt MA (1997) Mitotic spindle positioning in Saccharomyces cerevisiae is accomplished by antagonistically acting microtubule motor proteins. J Cell Biol 138: 1041–53

14. Craft Van De Weghe J, Harris JA, Kubo T, Witman GB & Lechtreck KF (2020) Diffusion rather than intraflagellar transport likely provides most of the tubulin required for axonemal assembly in Chlamydomonas. J Cell Sci 133

15. Drechsler H, Tan AN & Liakopoulos D (2015) Yeast GSK-3 kinase regulates astral microtubule function via phosphorylation of the microtubule-stabilizing kinesin Kip2. J Cell Sci 128: 3910–3921

16. Friel CT & Welburn JP (2018) Parts list for a microtubule depolymerising kinesin. Biochem Soc Trans 46: 1665–1672

17. Galbraith JA & Gallant PE (2000) Axonal transport of tubulin and actin. J Neurocytol 29: 889–911

18. Grava S & Philippsen P (2010) Dynamics of multiple nuclei in Ashbya gossypii hyphae depend on the control of cytoplasmic microtubules length by Bik1, Kip2, Kip3, and not on a capture/shrinkage mechanism. Mol Biol Cell 21: 3680–92

19. Hibbel A, Bogdanova A, Mahamdeh M, Jannasch A, Storch M, Schäffer E, Liakopoulos D & Howard J (2015) Kinesin Kip2 enhances microtubule growth in vitro through length-dependent feedback on polymerization and catastrophe. Elife 4: e10542

20. Hoffman PN & Lasek RJ (1980) Axonal transport of the cytoskeleton in regenerating motor neurons: constancy and change. Brain Res 202: 317–333

21. Howard J & Hyman A a (2007) Microtubule polymerases and depolymerases. Curr Opin Cell Biol 19: 31–5

22. Hunter AW & Wordeman L (2000) How motor proteins influence microtubule polymerization dynamics. J Cell Sci 113 Pt 24: 4379–89

23. Hyman A, Drechsel D, Kellogg D, Salser S, Sawin K, Steffen P, Wordeman L & Mitchison T (1991) Preparation of modified tubulins. In Methods in Enzymology pp 478–485.

24. Jaulin F & Kreitzer G (2010) KIF17 stabilizes microtubules and contributes to epithelial morphogenesis by acting at MT plus-ends with EB1 and APC. J Cell Biol 190: 443–60

25. Lawrence CJ, Malmberg RL, Muszynski MG & Dawe RK (2002) Maximum likelihood methods reveal conservation of function among closely related kinesin families. J Mol Evol 54: 42–53

26. Lee WL, Kaiser MA & Cooper JA (2005) The offloading model for dynein function. J Cell Biol 168: 201–207

27. Maekawa H, Usui T, Knop M & Schiebel E (2003) Yeast Cdk1 translocates to the plus end of cytoplasmic microtubules to regulate bud cortex interactions. EMBO J 22: 438–49

28. Markus SM, Punch JJ & Lee W-L (2009) Motor- and tail-dependent targeting of dynein to microtubule plus-ends and the cell cortex. Curr Biol 19: 196–205

29. Meier SM, Farcas AM, Kumar A, Ijavi M, Bill RT, Stelling J, Dufresne ER, Steinmetz MO & Barral Y (2023) Multivalency ensures persistence of a +TIP body at specialized microtubule ends. Nat Cell Biol 25: 56–67

30. Miesch J, Wimbish RT, Velluz MC & Aumeier C (2023) Phase separation of +TIP networks regulates microtubule dynamics. Proc Natl Acad Sci 120: 1–37

31. Miller RK, Heller KK, Frisèn L, Wallack DL, Loayza D, Gammie AE & Rose MD (1998) The kinesin-related proteins, Kip2p and Kip3p, function differently in nuclear migration in yeast. Mol Biol Cell 9: 2051–68

32. Moores CA, Perderiset M, Kappeler C, Kain S, Drummond D, Perkins SJ, Chelly J, Cross R, Houdusse A & Francis F (2006) Distinct roles of doublecortin modulating the microtubule cytoskeleton. EMBO J 25: 4448–4457

33. Raudaskoski M (2022) Kinesin Motors in the Filamentous Basidiomycetes in Light of the Schizophyllum commune Genome. J Fungi 8: 294

34. Roberts AJ, Goodman BS & Reck-Peterson SL (2014) Reconstitution of dynein transport to the microtubule plus end by kinesin. Elife 3: 1–16

35. Sardar HS, Luczak VG, Lopez MM, Lister BC & Gilbert SP (2010) Mitotic kinesin CENP-E promotes microtubule plus-end elongation. Curr Biol 20: 1648–53

36. Shrestha S, Hazelbaker M, Yount AL & Walczak CE (2018) Emerging Insights into the Function of Kinesin-8 Proteins in Microtubule Length Regulation. Biomolecules 9: 1

37. Steinberg G (2007) Preparing the way: fungal motors in microtubule organization. Trends Microbiol 15: 14–21

38. Su X, Ohi R & Pellman D (2012) Move in for the kill: motile microtubule regulators. Trends Cell Biol 22: 567–75

39. Uchida A, Alami NH & Brown A (2009) Tight Functional Coupling of Kinesin-1A and Dynein Motors in the Bidirectional Transport of Neurofilaments. Mol Biol Cell 20: 4997–5006

40. Wickstead B & Gull K (2006) A ‘holistic’ kinesin phylogeny reveals new kinesin families and predicts protein functions. Mol Biol Cell 17: 1734–43

41. Wu X, Xiang X & Hammer J a (2006) Motor proteins at the microtubule plus-end. Trends Cell Biol 16: 135–43

